# *In silico* APC/C substrate discovery reveals cell cycle degradation of chromatin regulators including UHRF1

**DOI:** 10.1101/2020.04.09.033621

**Authors:** Jennifer L. Kernan, Raquel C. Martinez-Chacin, Xianxi Wang, Rochelle L. Tiedemann, Thomas Bonacci, Rajarshi Choudhury, Derek L. Bolhuis, Jeffrey S. Damrauer, Feng Yan, Joseph S. Harrison, Michael Ben Major, Katherine Hoadley, Aussie Suzuki, Scott B. Rothbart, Nicholas G. Brown, Michael J. Emanuele

## Abstract

The Anaphase-Promoting Complex/Cyclosome (APC/C) is an E3 ubiquitin ligase and critical regulator of cell cycle progression. Despite its vital role, it has remained challenging to globally map APC/C substrates. By combining orthogonal features of known substrates, we predicted APC/C substrates *in silico*. This analysis identified many known substrates and suggested numerous candidates. Unexpectedly, chromatin regulatory proteins are enriched among putative substrates and we show that several chromatin proteins bind APC/C, oscillate during the cell cycle and are degraded following APC/C activation, consistent with being direct APC/C substrates. Additional analysis revealed detailed mechanisms of ubiquitylation for UHRF1, a key chromatin regulator involved in histone ubiquitylation and DNA methylation maintenance. Disrupting UHRF1 degradation at mitotic exit accelerates G1-phase cell cycle progression and perturbs global DNA methylation patterning in the genome. We conclude that APC/C coordinates crosstalk between cell cycle and chromatin regulatory proteins. This has potential consequences in normal cell physiology, where the chromatin environment changes depending on proliferative state, as well as in disease.

## Introduction

Regulated protein degradation is central to cell and organismal physiology and plays a particularly important role in proliferation. In eukaryotes, protein degradation is controlled largely by the ubiquitin (Ub) system. E3 Ub ligases provide substrate specificity and facilitate the transfer of Ub onto substrates. The formation of poly-Ub chains on substrates provides a signal that often targets substrates to the proteasome for degradation (1).

The Anaphase-Promoting Complex/Cyclosome (APC/C) is a 1.2 megadalton, multi-subunit E3 ligase and essential cell cycle regulator. APC/C utilizes two coactivators, Cdc20 and Cdh1, which directly bind substrates, recruiting them to the E3 complex (2). APC/C^Cdc20^ becomes active in mid-mitosis and promotes the metaphase to anaphase transition. APC/C^Cdh1^ becomes active in late mitosis and remains active until the end of G1, during which time it prevents S-phase entry (3). Thus, APC/C^Cdc20^ and APC/C^Cdh1^ play opposing roles, the former promoting cell cycle progression in mitosis and the latter inhibiting cell cycle progression in G1.

In addition to its role in normal cell cycles, APC/C dysfunction has been implicated in disease. Cdh1 is a haploinsufficient tumor suppressor in mice and cooperates with the retinoblastoma protein to restrain proliferation (4–8). Several oncogenic kinase cascades impinge on Cdh1 function, further supporting a role for APC/C^Cdh1^ in tumor suppression (9–11). In addition, the APC/C subunit Cdc27 is mutated in cancer and associated with aneuploidy (12). APC/C is also linked to inherited disorders that give a range of disease phenotypes, including microcephaly, cancer predisposition, and skeletal abnormalities (13,14).

Cdh1 and Cdc20 bind substrates through short, linear sequence motifs termed degrons. The most well-defined APC/C degron motifs are the KEN-box and D-box (15,16). In addition, binding of Cdc20 and Cdh1 to APC/C promotes a conformational change in the E3 that stimulates ligase activity (17). This results in substrate poly-ubiquitylation by its two cognate E2 enzymes. UBE2C/UbcH10 deposits the first Ub monomers onto substrates and forms short Ub chains, whereas UBE2S elongates poly-Ub chains (18–21).

Most known APC/C substrates are linked to cell cycle processes, including mitotic progression, spindle function and DNA replication. The paramount importance of APC/C in cell cycle and non-cell cycle processes, and its dysfunction in disease, highlight the importance of systematically defining substrates, whose regulation (or dysregulation), will likely contribute to proliferation and disease phenotypes. Nevertheless, barriers exist to the identification of APC/C substrates, as well as most other E3s. E3-substrate interactions are dynamic and binding often triggers substrate proteolysis. Additionally, the abundance of most substrates is low, and for APC/C, most targets are cell cycle regulated. Furthermore, since APC/C is a massive complex with many substrates, the relative binding stoichiometry to each individual substrate is low. Finally, degron sequences are short and occur vastly across proteomes, making it difficult to predict substrates.

We developed a simple *in silico* approach to identify potential APC/C targets. We took advantage of common features among known substrates, namely their transcriptional regulation during cell cycle and the presence of a degron motif. Super-imposing these features onto the proteome enriched for substrates and suggested previously undescribed targets.

This analysis revealed a role for APC/C in chromatin biology. We validate several substrates involved in chromatin dynamics, highlighting a previously underappreciated role for APC/C in chromatin regulation. We further define the mechanisms of ubiquitylation for UHRF1 (<u>U</u>biquitin-like with P<u>H</u>D and <u>R</u>ING <u>f</u>inger domains 1), a multivalent chromatin binding protein and itself an E3 ligase that can ubiquitylate histone H3 (22–25). UHRF1 plays an important role in DNA methylation and has been implicated in other DNA templated processes, including DNA repair (26–28). Additionally, UHRF1 is suggested to be an oncogene, whose expression correlates with high tumor grade and poor prognosis (29–31).

Altogether, these results reveal a role for APC/C-dependent UHRF1 degradation in cell cycle progression and shaping the DNA methylation landscape. More broadly, our data suggest that cell cycle regulated protein degradation helps organize the epigenetic landscape during proliferation. This suggests a potential mechanistic link contributing to changes in the chromatin landscape observed between proliferating and non-proliferating cells (32,33). We predict that altering APC/C function could promote changes in the histone and DNA modification landscape, and that these effects could contribute to the biochemical and phenotypic features of diseases, including cancer and neurological disorders.

## Results

### Identification of APC/C substrates

To identify human APC/C substrates, we first performed FLAG immunoprecipitations (IP) from HEK-293T cells expressing amino-terminal tagged FLAG-Cdh1 or an empty vector and analyzed precipitated proteins by mass spectrometry (Table S1). Several APC/C complex components and known substrates, including Rrm2, Kif11, Claspin, and cyclin A were enriched in Cdh1 pulldowns. Compared to a previously established dataset (34), we identified 15 out of 53 known substrates. However, hundreds of proteins were enriched over controls and many known substrates scored weakly, confounding our ability to prioritize candidates. For example, a single spectral count was observed for the substrate Kif22/KID (35,36).

We considered computationally identifying substrates based on features common among substrates. APC/C binds substrates most often through D- and KEN-box degron motifs. The minimal D-box motif (R-x-x-L) is present in most human proteins and insufficient as a prediction tool. The KEN-motif is found in approximately 10% of human proteins (2,206; Table S2), and several D-box regulated substrates also contain a KEN-motif, including Securin and Cdc6 (37,38). In addition, the gene expression of most APC/C substrates oscillates during the cell cycle (39). We cross-referenced the KEN-motif containing proteins against a set of 651 proteins whose mRNAs scored in at last two cell cycle mRNA profiling studies (40–43). Overlapping the 2,206 KEN-motif containing proteins with 651 transcriptionally controlled genes produced a set of 145 proteins, which represent known and putative APC/C substrates (Fig. 1A, Table S2).

**Fig. 1.**
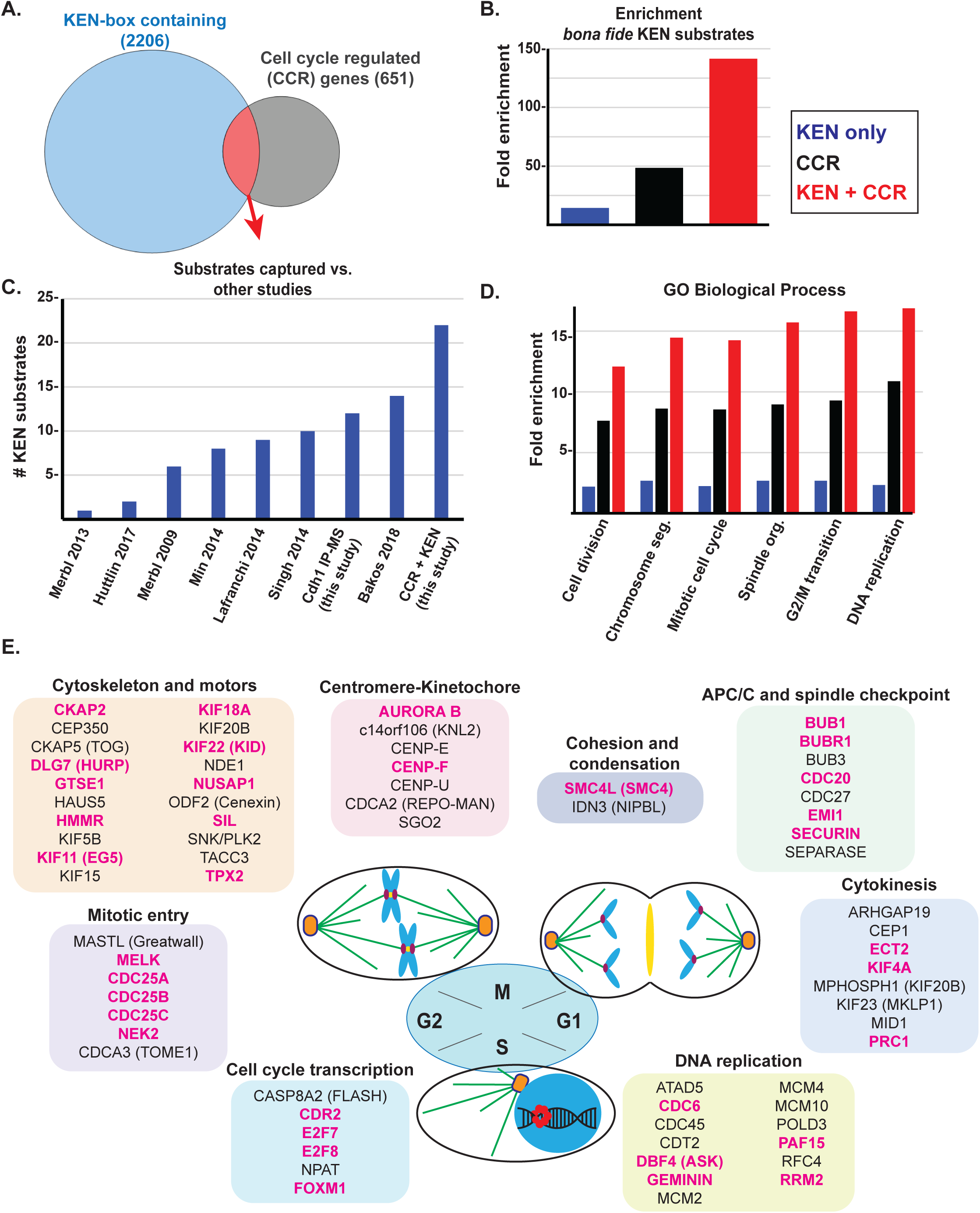
*In silico* analysis reveals a high confidence set of APC/C substrates involved in mitosis. (A) KEN-box containing human proteins were identified and cross-referenced against a set of 651 genes whose expression is cell cycle regulated based on multiple, independent studies. This revealed a set of 145 KEN-box containing proteins whose mRNA expression is cell cycle regulated. (B) Analysis of the enrichment of *bona fide* KEN-dependent substrates among these three datasets (blue- KEN box only set (2206); black- cell cycle regulated mRNAs (651); red- the overlapping set of 145 proteins) compared against a curated set of *bona fide*, KEN-dependent APC/C substrates (Davey and Morgan, Mol Cell, 2016). Enrichment was calculated based on the expected number of substrates which would be captured by chance based on the size of the dataset. (C) Analysis of putative substrates recovered in the indicated studies. (D) Gene ontology (GO) analysis for indicated studies (blue- KEN box only set (2206); black- cell cycle regulated mRNAs (651); red- the overlapping set of 145 proteins). (E) The set of 145 putative substrates was manually curated and analyzed for roles in various aspects of cell cycle progression. Seventy proteins, involved in cell cycle activities, are shown. The ones labelled in magenta signify that there is evidence in the literature of their regulation by APC/C. (Note that AURORA B, a mitotic kinase that phosphorylates histone H3, is listed here and in Figure 2A)

We compared our *in silico* analysis with two previously curated datasets, one containing 53 known APC/C targets (34), and a second containing 33 specifically KEN-dependent APC/C substrates (16). When compared to these lists of 53 and 33 substrates, our dataset captured 26 and 22 of them, respectively, the latter representing an enrichment of more than 140-fold, compared to what would be expected by chance (Fig. 1B). We also compared our data to other studies that identified APC/C substrates, interactors, proteins degraded at mitotic exit, or proteins ubiquitylated in mitosis (Table S3) (34,44–49). Our *in silico* analysis identified the most KEN-dependent substrates relative to these studies (Figure 1C; Table S3). When compared to the set of 53 substrates, which includes both D- and KEN-box dependent substrates, our dataset captured 26 out of 53 known substrates, despite not focusing on D-box substrates. Combining the *in silico* predictions with our Cdh1-pulldown proteomics data, we captured 31 out of 53 substrates.

Among the 145 computationally identified known and potential substrates, gene ontology (GO) analysis showed a strong enrichment for processes linked to various aspects of cell division (Fig. 1D). Manual curation demonstrated that nearly half of the proteins we identified (70 of 145) have well-established roles in cell cycle. These were sub-classified into the sub-categories cytoskeleton and motors, centromere-kinetochore, APC/C and spindle checkpoint, cytokinesis, mitotic entry, cell cycle transcription, cohesion and condensation, and DNA replication (Fig. 1E). Among these 70, 50% have literature evidence for regulation by APC/C, highlighting our enrichment for APC/C substrates (Fig. 1E; shown in magenta). All 145 proteins, their known function, sub-category, KEN-box sequence motif with flanking sequence, aliases, and citations describing regulation by APC/C are detailed in Table S2.

### Regulated degradation of chromatin factors

Unexpectedly, our dataset revealed several proteins involved in chromatin regulation (Fig. 2A) and an enrichment for GO processes related to chromatin (Fig. 2B). The dataset includes readers and writers of histone post-translational modifications, including the lysine acetyltransferases, PCAF/KAT2B and NCOA3/KAT13B, the lysine methyl-transferase MLL2/KMT2D, the chromatin reader and histone Ub ligase UHRF1, and the mitotic histone H3 kinase Aurora B (Fig. 2A and 1E). We identified proteins involved in chromatin assembly and structure, including: CHAF1B, a component of the CAF-1 nucleosome assembly complex; TTF2, a Swi2/Snf2 family member and DNA-dependent ATPase; KI-67, which prevents chromosome aggregation in mitosis and regulates histone post-translational modifications; and proteins associated with cohesion and condensation, including SMC4 and NIPBL (Fig. 1E). We also identified proteins involved in DNA damage repair.

**Fig. 2.**
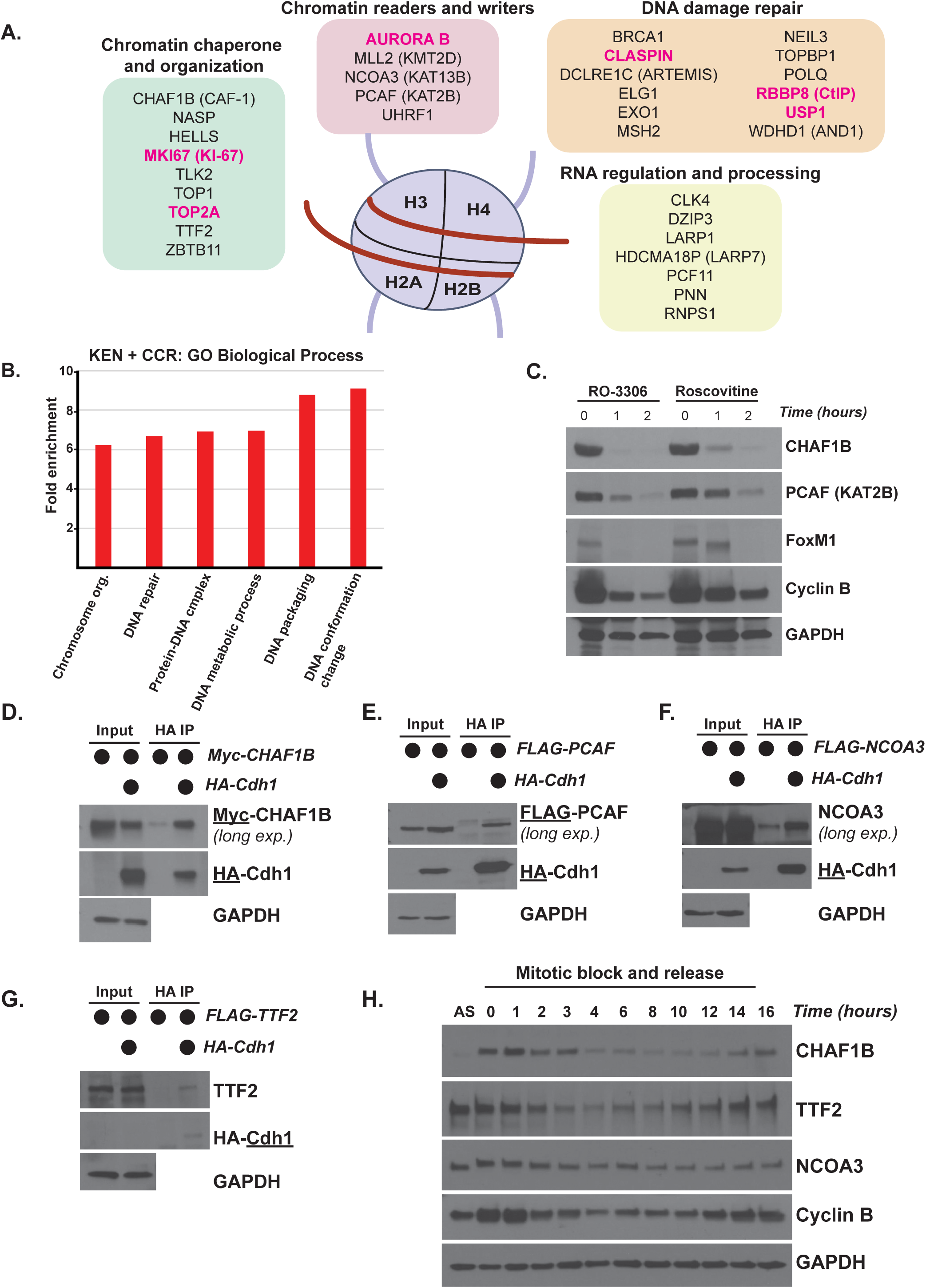
Putative APC/C substrates are enriched for roles in chromatin regulation. (A) The set of 145 known and putative APC/C substrates is enriched for proteins involved in various chromatin related process. This includes chromatin readers and writers, chaperones, RNA regulation and processing, DNA damage repair, and others. (Note that AURORA B, a mitotic kinase that phosphorylates histone H3, is listed here and in Figure 1E) (B) Gene ontology (GO) analysis of the overlapping KEN-box containing cell cycle regulated transcripts. This set is enriched for the indicated biological process, including DNA metabolism, protein-DNA complex assembly, DNA packaging, and DNA conformation. (C) APC/C activation assay to monitor substrate degradation. Following synchronization in mitosis, cells were washed one time and treated with CDK inhibitors to remove inhibitory phosphorylation marks that hinder the formation of APC/C^Cdh1^ needed for the M/G1 phase transition. Protein degradation was monitored by immunoblot. CHAF1B and PCAF are putative APC/C substrates, and FoxM1 and Cyclin B are known targets. (D) coIP of HA-Cdh1 with Myc-CHAF1B in transiently transfected 293T cells treated with proteasome inhibitors prior to harvesting. The underline indicates which protein or tag was blotted for in a particular panel (here and below).. Input equal to 1% of IP, here and below. (E) coIP of HA-Cdh1 with FLAG-PCAF in transiently transfected 293T cells treated with proteasome inhibitors prior to harvesting. (F) coIP of HA-Cdh1 with FLAG-NCOA3 in transiently transfected 293T cells treated with proteasome inhibitors prior to harvesting (G) coIP of HA-Cdh1 with FLAG-TTF2 in transiently transfected 293T cells treated with proteasome inhibitors prior to harvesting. (H) Mitotic shake-off of synchronized U2OS cells collected after release at the indicated timepoints. Immunoblotting for select endogenous proteins that are putative APC/C substrates or the positive control Cyclin B.

To validate potential substrates, we developed an *in vivo* APC/C activation assay that is amenable to analysis of endogenous or exogenously expressed proteins, and which is similar to approaches described elsewhere (50). U2OS cells were synchronized in mitosis with the microtubule poison nocodazole. After harvesting cells by mitotic shake-off, CDK1 was inactivated with either the CDK1-specific inhibitor RO-3306 or pan-CDK inhibitor Roscovitine, driving cells out of mitosis and triggering APC/C activation and destruction of substrates, including FoxM1, NUSAP1, and Cyclin B (Fig. 2C, Fig. S1) (51).

Using a combination of exogenous expression and endogenous protein analysis, we examined the levels of chromatin related proteins not previously shown to be APC/C substrates. Using this assay, there was a decrease in the levels of several writers of histone modifications, including UHRF1, PCAF, TTF2, and NCOA3 (Fig. 2C, S1A, S1B). We observed a decrease in the levels of the chromatin assembly factors NASP and CHAF1B as well as the RNA processing proteins LARP1 and LARP7 (Fig. 2C, S1A, S1B). All of these have been previously identified as ubiquitylated in proteomics studies by an unknown E3 ligase (52–56).

Since the role of APC/C in chromatin regulation is not well established, we focused our attention on the potential regulation of chromatin proteins by APC/C. We determined the ability of a subset to bind Cdh1 by coIP. CHAF1B, PCAF, NCOA3, and TTF2 interact with Cdh1 by coIP in 293T cells (Fig. 2D-2G). Accordingly, the levels of endogenous CHAF1B, TTF2, and NCOA3 oscillate during the cell cycle in U2OS, analyzed following a nocodazole-induced block in mitosis and then release into the cell cycle (Fig. 2H). PCAF levels did not decrease at mitotic exit in U2OS (Fig. S1C) but do decrease at mitotic exit in HeLa cells (Fig. S1C), suggesting a potentially complex regulation. Finally, we purified recombinant TTF2 and found that APC/C could trigger its ubiquitylation *in vitro* (Fig. S2). A table of all proteins tested in these assays and their validation is shown in Table S4. Taken together, this analysis uncovered new APC/C substrates and a role for APC/C in controlling chromatin regulators.

### UHRF1 regulation by APC/C^Cdh1^

To further understand the function of APC/C in chromatin biology, we pursued UHRF1, a key chromatin regulator that reads and writes histone modifications. UHRF1 associates with the DNA methyltransferase DNMT1 and is required for DNA methylation (26). UHRF1 has also been implicated in replisome assembly (57,58) and its phosphorylation oscillates during the cell cycle (59).

We examined UHRF1 protein levels following a mitotic block and release. Immunoblotting for UHRF1 and other cell cycle markers showed that UHRF1 protein levels decrease during mitotic exit in HeLa S3, HeLa, and U2OS cell lines (Fig. 3A, S3A-B). In each cell line, UHRF1 levels remain low in G1 and then re-accumulate starting around G1/S, based on the expression of other cell cycle markers, including cyclin E and cyclin A, and then further increasing throughout the subsequent G2/M phase.

**Fig. 3.**
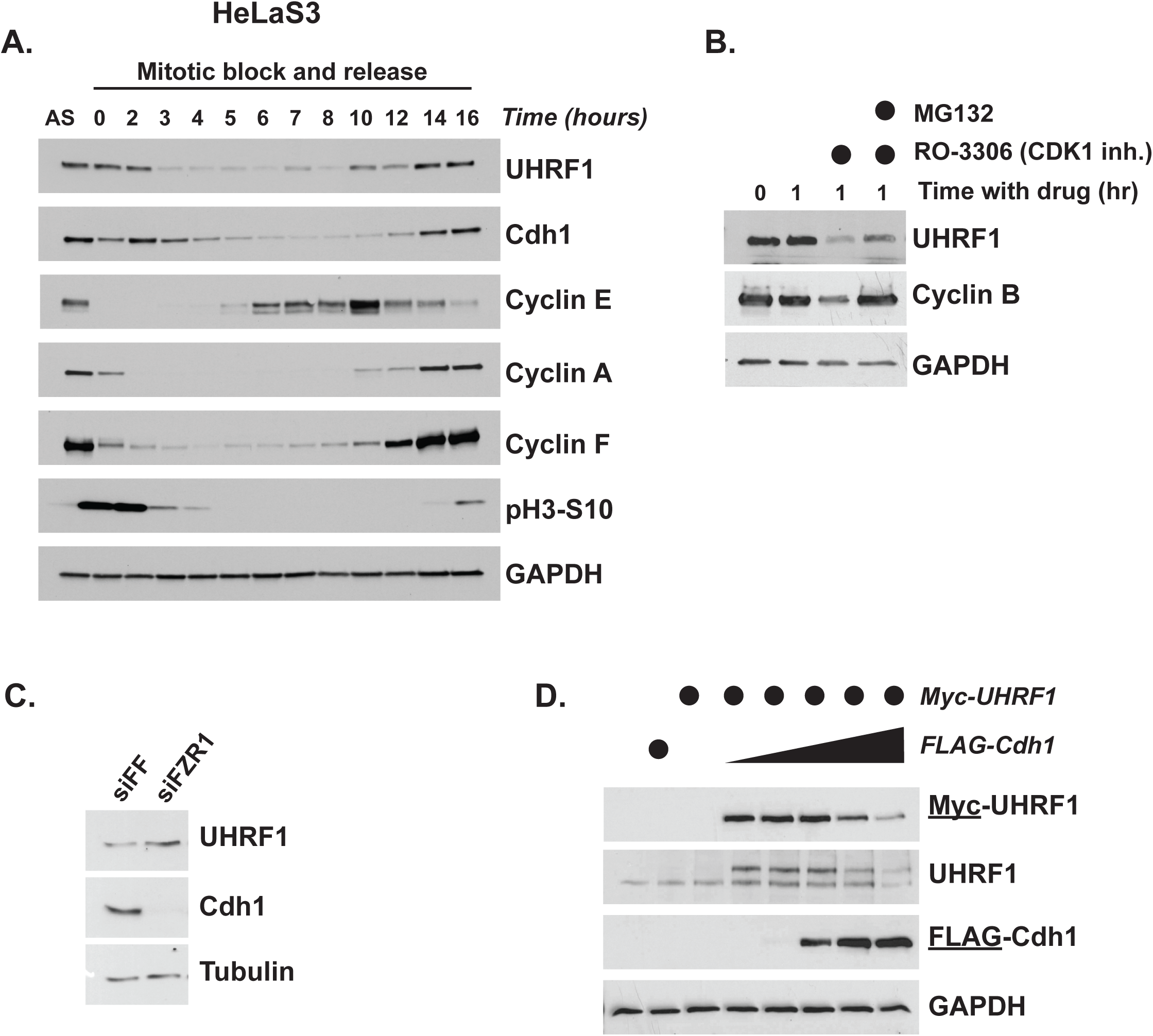
UHRF1 levels are controlled by APC/C^Cdh1^. (A) HeLa S3 cells were synchronized in mitosis and released into the cell cycle. Timepoints were taken at the indicated time points and analyzed by immunoblot. (B) U2OS cells were synchronized in prometaphase with 250ng/mL nocodazole for 16hr prior to mitotic shake-off. Cells were released into fresh media containing 10µM RO-3306 CDK inhibitor (used as described in Fig. 2C) with or without addition of 20µM of proteasomal inhibitor MG-132 and harvested 1hr later. Cyclin B is a positive control for a known APC/C substrate that is degraded at mitotic exit. (C) HCT116 cells were transfected with siRNA targeting Cdh1 (Fzr1 mRNA) or firefly luciferase as a control and harvested after 24 hr for immunoblotting. (D) Myc-UHRF1 was transiently expressed in 293T cells with increasing concentrations of FLAG-Cdh1 for 24hr before analysis by immunoblot.

We performed several assays to assess whether UHRF1 is regulated by APC/C. We analyzed UHRF1 in the aforementioned *in vivo* APC/C activation assay. U2OS cells were arrested in mitosis and then treated with RO-3306. We observed a decrease in UHRF1 that was partially mitigated by the proteasome inhibitor, MG-132, indicating that the reduction is dependent on the proteasome (Fig. 3B). In addition, transient siRNA depletion of Cdh1 (Fzr1 mRNA transcript) augmented UHRF1 protein levels (Fig. 3C). Conversely, ectopic expression of increasing concentrations of FLAG-Cdh1 led to a dose-dependent decrease in both exogenous and endogenous UHRF1 protein levels (Fig. 3D). We examined UHRF1 levels in cells that were first synchronized in G1 by a mitotic block and release, and then treated with the pharmacological APC/C inhibitor proTAME for 90 minutes (Fig. S3C). This led to an increase in UHRF1 levels. Together, these data suggest that APC/C controls UHRF1 *in vivo*.

### UHRF1 ubiquitylation by APC/C^Cdh1^

UHRF1 is a multi-domain protein (Fig. 4A) that exhibits multivalent binding with chromatin through histone and DNA binding domains (24,60,61). Additionally, UHRF1 is a RING domain E3 that ubiquitylates histone H3 (22,23,25). To determine whether UHRF1 is a direct APC/C^Cdh1^ substrate, we tested its binding to Cdh1 by expressing HA-Cdh1 and Myc-UHRF1 in 293T cells. Cells were treated with the proteasome inhibitor MG-132 prior to harvesting to prevent UHRF1 degradation. Myc-UHRF1 was enriched in the HA-Cdh1 pull-down, and HA-Cdh1 was enriched in the Myc-UHRF1 pull-down (Fig. 4B, 4C).

**Fig. 4.**
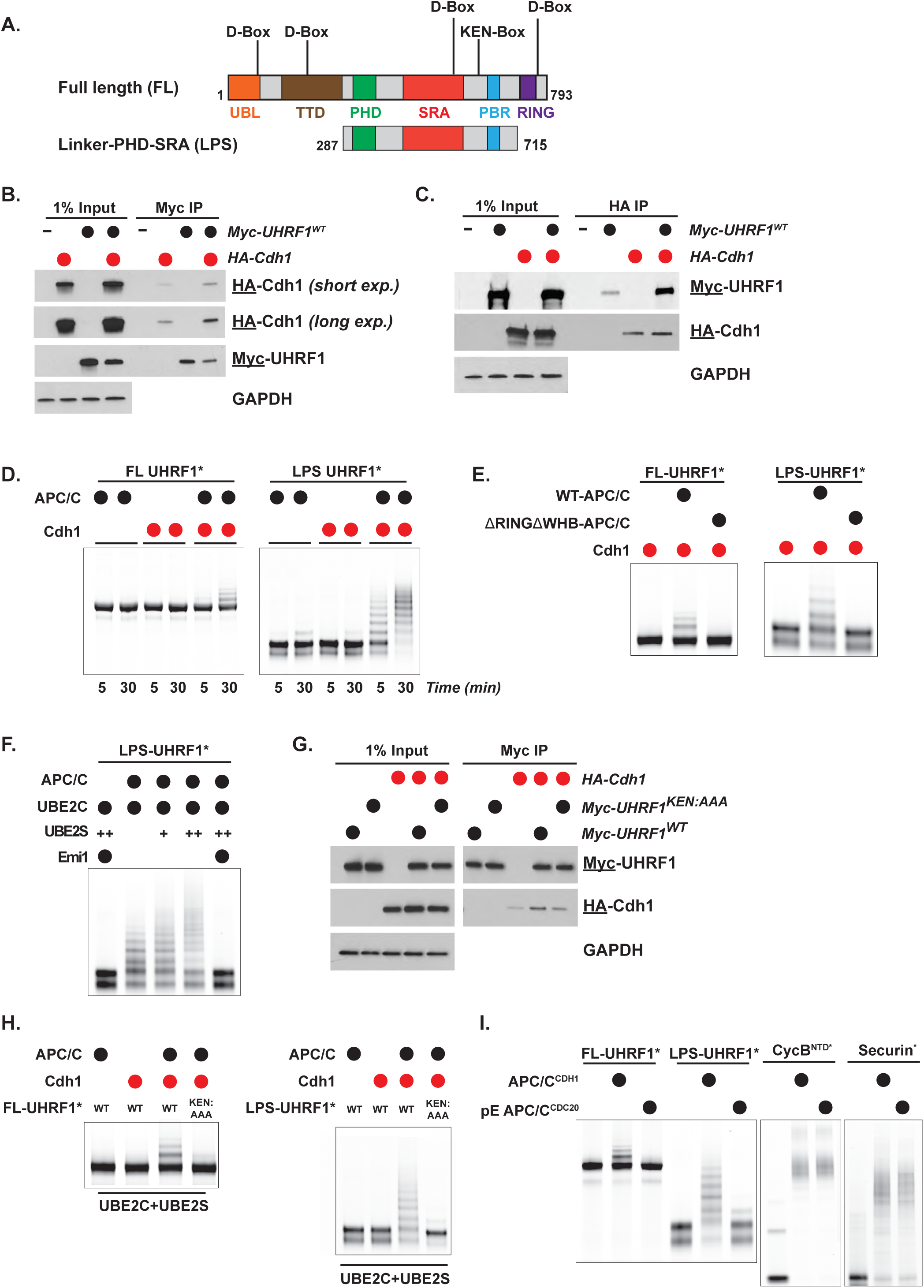
UHRF1 binding and ubiquitylation by APC/C^Cdh1^ depends on KEN degron. (A) Schematic of UHRF1 domain structure with location of KEN degron in both full-length (FL) and truncated LPS UHRF1. (B) coIP of HA-Cdh1 with Myc-UHRF1 in transiently transfected 293T cells treated with proteasome inhibitors prior to harvesting and α-Myc IP. Input equal to 1% of IP, here and below. (C) coIP of Myc-UHRF1 with HA-Cdh1 in transiently transfected 293T cells treated with proteasome inhibitors prior to harvesting and α-HA IP. (D) Ubiquitylation reactions with APC/C^Cdh1^, UBE2C, FL UHRF1* or LPS UHRF1*, and wild-type ubiquitin. UHRF1 was detected by fluorescence scanning (* indicates fluorescently labeled protein). (E) Ubiquitylation reactions similar as in (D) but using two variants of APC/C: WT and catalytically dead APC/C^ΔRINGΔWHB^, a version of APC/C that can neither recruit nor activate its E2, UBE2C. UHRF1 was detected by fluorescence scanning. Samples were collected at 30 min. (F) Representative *in vitro* ubiquitylation reactions showing UBE2S-dependent chain elongation reactions of LPS UHRF1*. Titration of UBE2S: 0 μM, 0.1 μM (+), 0.5 μM (++). The addition of Emi1 completely inhibited the reaction. UHRF1 was detected by fluorescence scanning. Samples were collected at 30 min. (G) coIP of HA-Cdh1 with Myc-UHRF1^WT^ or Myc-UHRF1^KEN:AAA^ in transiently transfected 293T cells treated with proteasome inhibitors prior to harvesting and α-Myc IP. (H) Polyubiquitylation reactions of FL-UHRF1* and LPS-UHRF1* by APC/C^Cdh1^, UBE2C, and UBE2S. UHRF1 ubiquitylation by APC/C^Cdh1^ is dependent on the KEN degron motif (lane 4 in both gels). UHRF1 was detected by fluorescence scanning. Samples were collected at 30 min. (I) Dependence of UHRF1 ubiquitylation on phosphorylation state of the APC/C (referred to as pE-APC/C) and subsequent coactivator recruitment. The well-established APC/C substrates, CycB^NTD^* and Securin*, are ubiquitylated by either APC/C^Cdc20^ or APC/C^Cdh1^, whereas UHRF1 is only ubiquitylated by APC/C^Cdh1^. Reactions were run in parallel. Collections taken at 1hr (for FL and LPS UHRF1*) and 30 min (for CycB^NTD^* and Securin*). Ubiquitylated proteins were detected by fluorescence scanning.

Next, we purified and fluorescently labeled recombinant, bacterially expressed, full-length (FL) UHRF1 (FL-UHRF1*, where the * denotes fluorescently labelled protein). We found that FL-UHRF1* was ubiquitylated in an APC/C- and Cdh1-dependent manner using an entirely *in vitro* recombinant system (Fig. 4D). Multiple, high molecular weight ubiquitylated forms are observed using either wild-type Ub or methylated-Ub, the latter of which cannot form poly-Ub chains. This indicates that APC/C ubiquitylates multiple lysines in UHRF1 (Fig. 4D, S4A-B).

Since UHRF1 can auto-ubiquitylate itself through its RING domain, we confirmed that its ubiquitylation is APC/C dependent. First, we purified a version of APC/C selectively missing the APC2 WHB domain and the APC11 RING domain, which are required to recruit its initiating E2 UBE2C (designated ΔRINGΔWHB) (62,63). This version of APC/C was unable to ubiquitylate UHRF1 (Fig. 4E).

Next, we purified and fluorescently labeled a truncated version of UHRF1 that contains the Linker, PHD, and SRA domains (termed LPS), spanning amino acids 287-715 (Fig. 4A). The LPS fragment omits three potential APC/C D-box degron motifs, as well as the RING domain, precluding auto-ubiquitylation. A D-box motif remains in the highly structured SRA domain but is unlikely to be accessible as a degron motif (64).

Significantly, LPS-UHRF1* is more robustly ubiquitylated in an APC/C- and Cdh1-dependent manner compared to FL-UHRF1* (Fig. 4D-E). Moreover, UHRF1 ubiquitylation is fully inhibited by the APC/C inhibitor Emi1 (Fig. 4F). Ubiquitylation of UHRF1 is initiated by APC/C^Cdh1^-UBE2C while APC/C^Cdh1^-UBE2S elongates Ub chains, indicating that UHRF1 ubiquitylation is similar to that of other substrates tested in this *in vitro* system (Fig. 4F). We conclude that UHRF1 is a *bona fide* APC/C substrate.

The ubiquitylation of truncated LPS-UHRF1* (Fig. 4D, 4E, 4F) strongly suggests the importance of the KEN-motif, located in an unstructured region at amino acids 622-624 (Fig. 4A). Alanine substitutions were introduced into the KEN sequence (UHRF1^KEN:AAA^). The KEN mutant version (Myc-UHRF1^KEN:AAA^) showed reduced, although not completely abolished, binding to HA-Cdh1 by coIP, compared to Myc-UHRF1^WT^ (Fig 4G). Additionally, the KEN mutant versions of FL-UHRF1* and LPS-UHRF1* were completely resistant to ubiquitylation by APC/C (Fig. 4H). We conclude that UHRF1 ubiquitylation by APC/C^Cdh1^ is dependent on its KEN-box motif.

APC/C substrates are recruited by Cdc20 and Cdh1, and many substrates can be controlled by both. To test if UHRF1 is controlled by APC/C^Cdc20^, in addition to APC/C^Cdh1^, we used a phosphomimetic version of APC/C (termed pE-APC/C) that can utilize either Cdc20 or Cdh1, since Cdc20 cannot bind to unphosphorylated APC/C (62). Surprisingly, unlike other, well-established APC/C substrates, including Cyclin B (CycB^NTD^, amino acids 1-95) and Securin, the FL-UHRF1* and LPS-UHRF1* were ubiquitylated by APC/C^Cdh1^ but not by APC/C^Cdc20^ (Fig. 4I, S4C).

We transiently expressed FLAG-Cdh1 in HEK-293T cells in combination with either Myc-UHRF1^WT^ or mutant versions harboring alanine mutations in either the KEN-box (Myc-UHRF1^KEN:AAA^) or the fourth D-box motif (Myc-UHRF1^D4^). Ectopic FLAG-Cdh1 overexpression triggers the degradation of Myc-UHRF1^WT^ and Myc-UHRF1^D4^, whereas Myc-UHRF1^KEN^ is resistant to degradation (Fig. 5A), further supporting the importance of the KEN-motif in UHRF1 degradation.

**Fig. 5.**
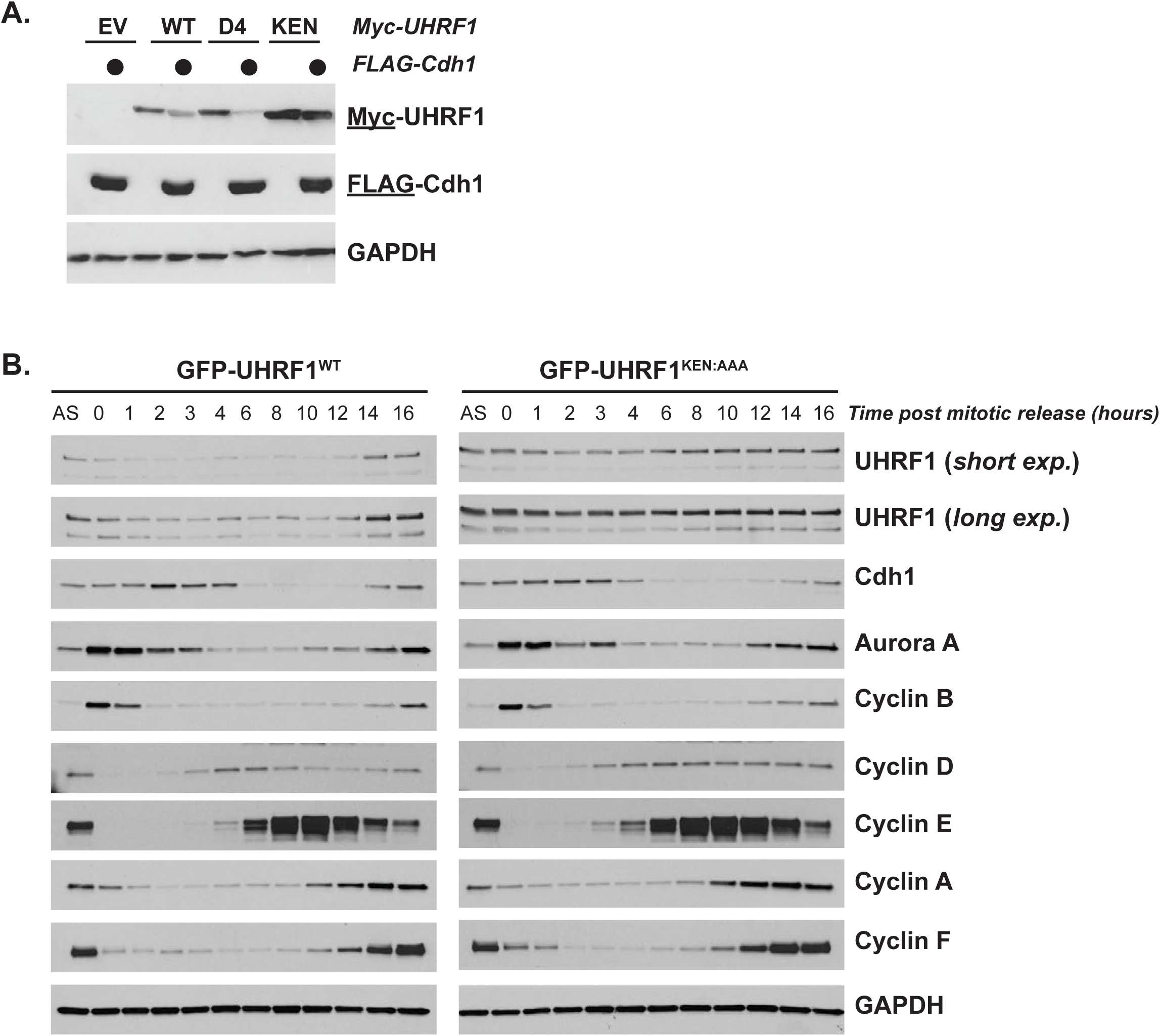
UHRF1 non-degradable mutant protein is stable at mitotic exit. (A) Myc-UHRF1^WT^ or mutant versions harboring alanine substitutions in either its KEN-box (KEN) or the fourth putative D-box motif (D4) (see Fig 4A for location of sequences) were transiently expressed in 293T cells with or without FLAG-Cdh1 for 24hr before analysis by immunoblot. (B) HeLa S3 stably expressing GFP-UHRF1^WT^ or GFP-UHRF1^KEN:AAA^ were synchronized in mitosis, released into the cell cycle, and collected for immunoblot analysis at the indicated timepoints.

Next, we generated cell lines constitutively expressing GFP-tagged UHRF1^WT^ or UHRF1^KEN:AAA^ using lentiviral transduction and examined UHRF1 stability upon mitotic exit. Exogenous UHRF1 levels were only moderately overexpressed compared to endogenous levels (Fig. 5B). Following synchronization with nocodazole, GFP-UHRF1^WT^ levels decrease at mitotic exit. Conversely, GFP-UHRF1^KEN:AAA^ levels remain stable through mitotic exit and G1 phase (Fig. 5B). Cells expressing GFP-UHRF1^KEN:AAA^ exit mitosis normally based on immunoblotting for the APC/C substrates cyclin A, cyclin B, cyclin F and Aurora A, which are degraded with normal kinetics (Fig. 5B). Thus, the KEN box regulates UHRF1 ubiquitylation *in vitro* and degradation *in vivo*. In addition, the mild over-expression of UHRF1 in these cells does not affect overall APC/C activity.

### UHRF1 degradation and cell cycle progression

Since many APC/C substrates are linked to proliferative control, we examined the contribution of UHRF1, and its degradation by APC/C, to cell cycle. Consistent with prior reports, UHRF1 depletion increased the fraction of cells in G1-phase ((65); data not shown). To further investigate the role of UHRF1 in cell cycle, we examined mitotic cells following UHRF1 depletion. We observed an approximately three-fold increase in cells with mis-aligned chromosomes in metaphase and anaphase in UHRF1 depleted cells using two independent siRNA oligonucleotides (Fig. S5A). Surprisingly, there was no statistically significant difference in the overall percent of mitotic cells.

To determine the role of UHRF1 degradation in cell cycle, we examined cell cycle markers in cells expressing UHRF1^WT^ or UHRF1^KEN:AAA^. In Hela cells traversing the cell cycle after synchronization at G1/S, following a double thymidine block and release, we found that the GFP-UHRF1^KEN:AAA^ cells contain more of the G1/S regulator cyclin E (Fig. S6A). This was also evident in cells that had been synchronized in mitosis and released into G1 (Fig. 5B). This suggested that an inability to degrade UHRF1 in G1 alters cyclin E expression, a key driver of S-phase entry.

These data suggested that UHRF1 might promote progression into S-phase and that a failure to degrade UHRF1 could shorten the duration of G1. To better address this possibility, we depleted endogenous UHRF1 with an shRNA targeting the UHRF1 3’UTR (66). Cells expressing GFP-UHRF1^WT^ or GFP-UHRF1^KEN:AAA^ were synchronized in mitosis, released into the cell cycle, and analyzed by immunoblot. Several markers of S-phase entry accumulate early in cells expressing GFP-UHRF1^KEN:AAA^ compared to GFP-UHRF1^WT^. Both cyclin E and the G1/S transcription factor E2F1 are elevated at early time points following release from mitosis (Fig. 6A). Elevated levels of cyclin E and E2F1 are evident in asynchronous RPE1-hTRET cells, and to a lesser extent in asynchronous HeLa S3 cells, where cell cycle transcription is perturbed due to HPV oncoproteins (Fig. S6B, S6C).

**Fig. 6.**
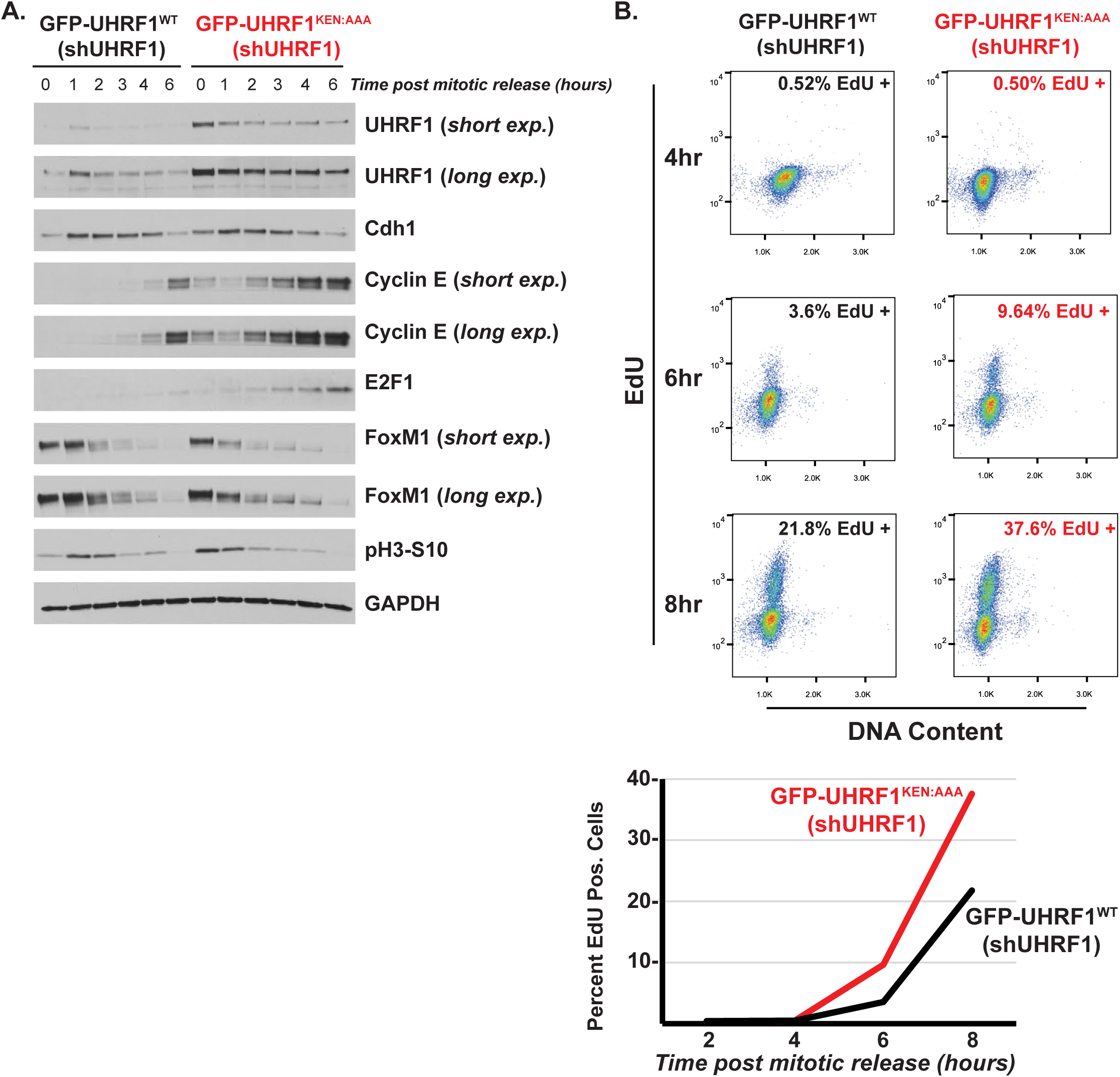
UHRF1 degradation restrains S-phase entry. (A) HeLa S3 stably expressing GFP-UHRF1^WT^ or GFP-UHRF1^KEN:AAA^ along with 3’UTR targeting shUHRF1 were synchronized in mitosis as described previously, released into the cell cycle, and collected for immunoblot analysis at the indicated timepoints, probing for cell cycle proteins as shown. (B) HeLa S3 stably expressing GFP-UHRF1^WT^ or GFP-UHRF1^KEN:AAA^ along with 3’UTR targeting shUHRF1 were synchronized in mitosis, released into the cell cycle, and pulsed with 10µM EdU for thirty minutes prior to harvest and analysis by flow cytometry. A representative experiment (n=3) is shown.

To analyze G1 duration, cells were release from a mitotic block and pulsed with EdU prior to harvesting for flow cytometry, to determine the percent of cells that were in S-phase. GFP-UHRF1^KEN:AAA^ expressing cells begin S-phase earlier than control cells (Fig. 6B). Six hours after release into the cell cycle, 3.6% of control cells had entered S-phase, whereas 9.6% of GFP-UHRF1^KEN:AAA^ expressing cells had started S-phase. Thus, a failure to degrade UHRF1 accelerates G1, indicating a key role for UHRF1 destruction in determining timing between the end of mitosis and start of DNA synthesis.

### UHRF1 degradation and DNA methylation homeostasis

UHRF1 is required for DNA methylation maintenance (26). To determine if stabilizing UHRF1 in G1 affects DNA methylation, we performed base-resolution DNA methylation analysis at approximately 850,000 unique human CpG loci spanning all genomic annotations and regulatory regions using the Infinium MethylationEPIC BeadChip (EPIC arrays) (67,68). We compared parental U2OS cells and those expressing GFP-UHRF1^WT^ or GFP-UHRF1^KEN:AAA^. Considering all probes, DNA methylation changes between parental, GFP-UHRF1^WT^, and GFP-UHRF1^KEN:AAA^ were insignificant (Fig. 7A). However, multidimensional scaling (MDS) of the top 50,000 variable CpG probes among all samples/replicates (agnostic of sample group) clustered experimental conditions (Fig. 7B), indicating a unique and reproducible profile of methylation patterning.

**Fig. 7.**
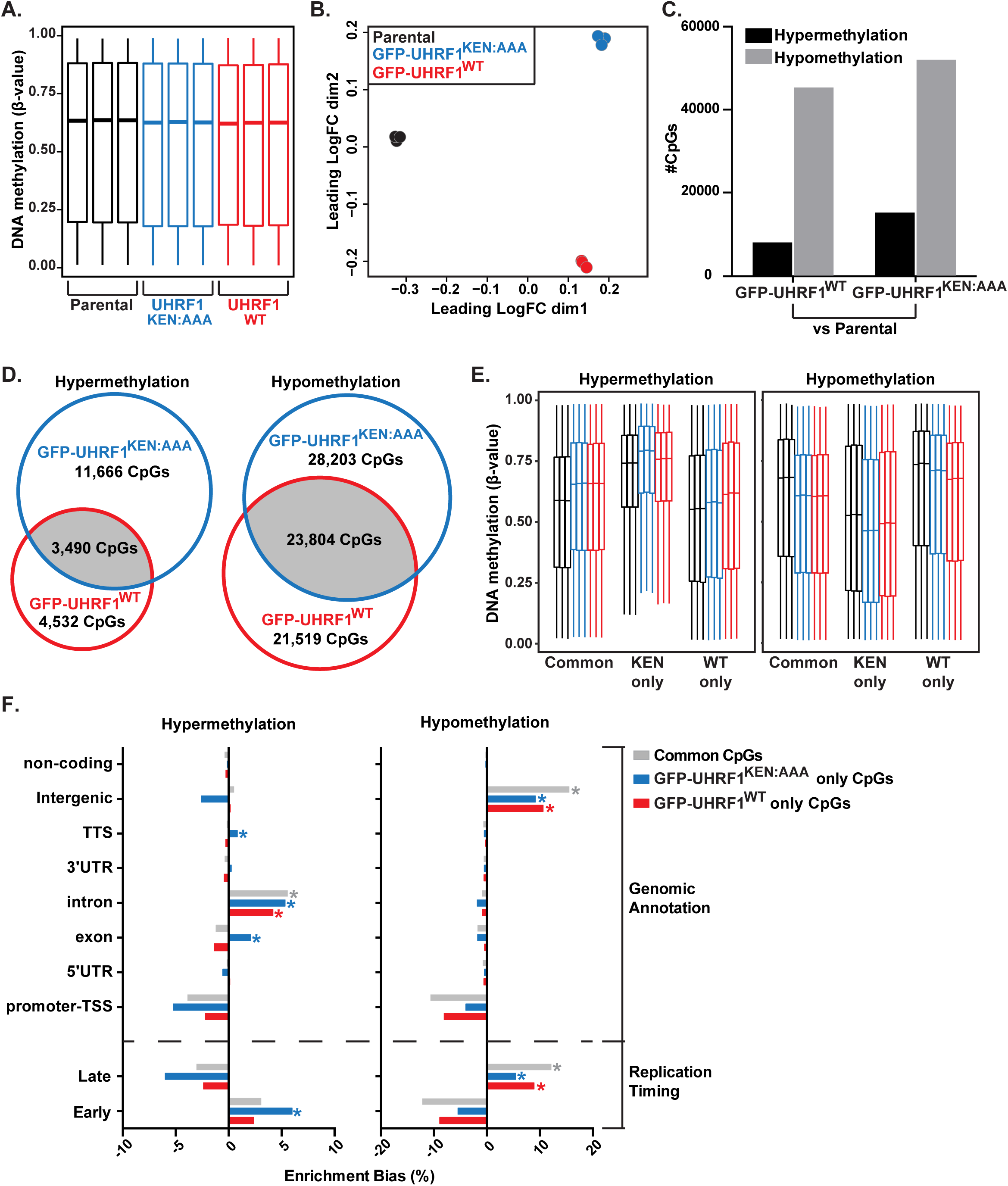
A non-degradable form of UHRF1 induces DNA hypermethylation of gene bodies and early replicating regions of the genome. (A) Global DNA methylation analysis for Parental U2OS and U2OS cells overexpressing GFP-UHRF1^WT^ or GFP-UHRF1^KEN:AAA^ with the Infinium MethylationEPIC BeadChip (Illumina) platform. Each sample group is represented in biological triplicate. All CpG probes that passed quality control analysis (n = 724,622 CpGs) are plotted as β-values population averages from 0 (fully unmethylated) to 1 (fully methylated). The midlines of each box plot represent the median DNA methylation value for all CpG probes in a sample. (B) Multidimensional scaling (MDS) of the top 50,000 variable CpG probes among samples. (C) Number of CpG probes that were differentially hypermethylated or hypomethylated in the GFP-UHRF1^WT^ and GFP-UHRF1^KEN:AAA^ groups relative to the Parental samples adjusted p-value <u>≤</u> 0.05). (D) Overlap analysis of significantly hypermethylated (left) or hypomethylated (right) CpG probes between GFP-UHRF1^WT^ and GFP-UHRF1^KEN:AAA^ sample groups. (E) DNA methylation levels of significantly hypermethylated (left) or hypomethylated (right) probes from (D) that are common between GFP-UHRF1^WT^ and GFP-UHRF1^KEN:AAA^ sample groups, unique to GFP-UHRF1^KEN:AAA^ (KEN only), or unique to GFP-UHRF1^WT^ (WT only). Color code from Fig. 7A applies. Outliers removed to simplify visualization. (F) Enrichment bias analysis of significantly hypermethylated (left) or hypomethylated (right) CpG probes among genomic annotations and U2OS replication timing data. *p-value <u>≤</u> 1E-300 for positive enrichment of the feature by hypergeometric testing.

We queried the GFP-UHRF1^WT^ and GFP-UHRF1^KEN:AAA^ samples for differentially methylated CpGs relative to the parental controls. Consistent with a previous report (29), expression of GFP-UHRF1^WT^ and GFP-UHRF1^KEN:AAA^ induced a comparable number of hypomethylation events (Fig. 7C). Altrenatively, GFP-UHRF1^KEN:AAA^ induced approximately two-fold more hypermethylated CpGs compared to GFP-UHRF1^WT^ (Fig. 7C). Analysis of differentially methylated CpG probes between GFP-UHRF1^WT^ and GFP-UHRF1^KEN:AAA^ revealed a 32% overlap in hypomethylated probes and a 17% overlap in hypermethylated probes (Fig. 7D). Significantly, hypermethylated CpG probes in the GFP-UHRF1^KEN:AAA^ expressing cells were 2.5-fold more abundant compared to GFP-UHRF1^WT^, despite no significant change in hypomethylated CpG probes. Thus, the non-degradable form of UHRF1 induces site-specific DNA hypermethylation (Fig. 7D).

The CpGs that were hypermethylated in GFP-UHRF1^KEN:AAA^ -expressing cells started with a higher methylation level than other categories and gained methylation due to expression of non-degradable mutant (Fig. 7E). Enrichment analysis of the differentially methylated CpGs revealed that gene body annotations, including exons, introns, and transcription termination sites (TTS), were positively enriched for hypermethylation in GFP-UHRF1^KEN:AAA^ -expressing cells (Fig. 7F, left panel). We next queried enrichment of differential methylation events in regions of early and late replication (69). Hypermethylation events in GFP-UHRF1^KEN:AAA^, but not GFP-UHRF1^WT^, were positively enriched in early replicating regions of the genome, while hypomethylation events by both GFP-UHRF1^WT^ and GFP-UHRF1^KEN:AAA^ (alone or shared in common) were enriched in late replicating DNA (Fig. 7F). The enrichment of these hypermethylated features was consistent with known DNA methylation patterns that occur across gene bodies and early replicating DNA (Fig. 7E), as CpG loci in these regions typically demonstrate high levels of methylation (70,71). Taken together, these results demonstrate that expression of non-degradable UHRF1 enhances methylation at gene-rich, early replicating regions of the genome.

## Discussion

### Identification of new E3 ligase substrates

APC/C is a core component of the cell cycle oscillator and mounting evidence points to its dysfunction in cancer and neurological disease. Here we provide a comprehensive, unencumbered, annotated list of known and candidate APC/C substrates. Our data highlights the importance of APC/C in various aspects of proliferative control and points to its potentially broader impact on unanticipated cellular processes, including chromatin organization.

Identifying E3 substrates remains technically challenging. Since E3-substrate interactions exhibit low stoichiometry, mapping substrates by defining interactors is difficult. In addition, Ub ligase substrates are often in low abundance. APC/C is inhibited throughout the cell cycle by myriad mechanisms (72) and the time when it binds substrates coincides with when targets are being degraded and their abundance is lowest. This complicates many proteomics-based approaches. Alternative techniques for identifying E3 ligase substrates, including Global Protein Stability Profiling (GPS) and *in vitro* expression cloning, circumvent these challenges by measuring changes in substrate stability using fluorescent reporters or metabolic labeling with radioisotopes. These represent powerful tools for mapping E3 substrates (56,73). However, both approaches are laborious and time intensive, require significant technical expertise, and depend on gene expression libraries, which are neither complete nor available to most laboratories.

We bypass these challenges using a simple *in silico* approach based on publicly available information, which is simple, inexpensive, and easily repeated with different variables. While our approach shares some similarities with previous approaches, it improves upon those in its simplicity, expanded use of multiple cell cycle mRNA datasets, and inclusion of a degron motif in the search criteria (35,39,74). Its success stems from the use of orthogonal filtering criteria, that is, unlinked features between mRNA and proteins. We predict that similar uses of unrelated properties could be leveraged for mapping targets of other enzymes, such as kinases, where defining substrates has proven similarly challenging. It is notable that degron sequences remain unknown for most Ub ligases, highlighting the importance of mechanistic studies in enabling systems-level discoveries.

### Involvement of APC/C in chromatin regulation

Determining the enzymes and substrates in kinase signaling cascades has been instrumental in determining proliferative controls in normal cells, their responses to stress and damage, and disease phenotypes and treatments. Relatedly, decoding Ub signaling pathways involved in proliferation is likely to provide insight into enzyme function in normal cell physiology as well as in disease.

A major finding of this work is that numerous chromatin regulators are controlled temporally during proliferation by APC/C. Impairing the degradation of one such substrate, UHRF1, altered the timing of cell cycle events and changed global patterns of DNA methylation. Since numerous chromatin regulators are controlled by APC/C, we anticipate widespread, pleiotropic effects on chromatin in cells where APC/C activity is impaired, either physiologically or pathologically.

Our observations raise the possibility that dysregulation of the cell cycle machinery, as is seen in diseases such as cancer, could alter the chromatin environment. The discovery that many chromatin regulators are mutated in cancer, a disease of uncontrolled proliferation, together with our data, imply a bidirectional relationship between the chromatin landscape and the cell cycle oscillator. Consistent with the notion that dysregulation of APC/C controlled proteins could play important roles in determining the chromatin environment in disease, the mRNA expression of our 145 known and putative substrates strongly predict breast cancer aneuploidies and copy number variations (Fig. S7). This observation is not due solely to the selection of specific breast cancer subtypes, since our gene signature is elevated in multiple breast cancer subtypes. Interestingly, the expression of this signature correlates with the CIN70 signature, which was previously developed based on gene expression in chromosomally unstable cancers (75). We observed an extraordinary correlation between the CIN70 and our 145 gene signature in breast cancer (Fig. S7). This is remarkable since our signature was generated completely independent of gene expression in cancer and was instead derived, in part, by short sequence motifs on proteins.

APC/C^Cdh1^, but not APC/C^Cdc20^, ubiquitylates UHRF1. This is notable because the Cdh1-bound form of APC/C is active both G1 and quiescent cells and is critical for restraining S-phase entry. Our findings suggest that impaired UHRF1 degradation promotes a premature G1/S transition. We propose that the proper degradation of UHRF1, and other chromatin regulators, serves to integrate growth factor dependent proliferative decisions with the chromatin regulatory environment. This could help explain the complex chromatin rearrangements observed in quiescent cells, where APC/C^Cdh1^ is active (32,33,76). Further, APC/C controls key cell cycle transcriptional regulators, including the G2/M transcription factor FoxM1 and the repressor E2F proteins, E2F7 and E2F8 (77,78). Thus, our data point to a higher order role regulatory role for APC/C in gene regulation, by controlling transcription factors (i.e. FoxM1), transcriptional repressors (i.e. E2F7, E2F8,) and chromatin modifiers.

Aberrant DNA methylation is a hallmark of cancer (79). UHRF1 promotes DNA methylation maintenance, and too much or too little UHRF1 expression is detrimental to methylation stasis (26,29). It is interesting to speculate that the redistribution of DNA methylation in disease could be caused, in part, by the aberrant stabilization of UHRF1, resulting from APC/C^Cdh1^ inactivation. It will be important, in the future, to determine if oncogene activation acts through the APC/C to re-organize the chromatin landscape. Furthermore, determining ubiquitin ligase substrates, like UHRF1, that might be dysregulated in pathological settings via altered degradative mechanisms could suggest therapeutic strategies to reverse their effects.

## Materials and Methods

### Computational identification of putative APC/C substrates

Human proteins containing a KEN-box sequence (amino acid sequence K-E-N) were identified using the “Find a Sequence Match” feature on the Scansite web search platform (currently https://scansite4.mit.edu/4.0/#home). Proteins with cell cycle regulated mRNA were curated from four independent cell cycle transcriptional studies (42,43,80,81). The genes which scored in two or more of these screens was previously compiled in the supplemental data of Grant et al., 2013. Gene and protein name conversions were performed using the DAVID online tool (https://david.ncifcrf.gov/conversion.jsp). The overlapping set 145 proteins, which contain a KEN sequence and exhibit oscillating cell cycle regulated mRNA expression, were identified. For all 145 proteins, we manually curated information on their alias, function, sequence flanking the KEN motif, and evidence for regulation by APC/C from various online databases and repositories, including UNIPROT, Pubmed, and Genecards.

The set of 33 well-validated, KEN-containing, human APC/C substrates was derived from (16). Our own FLAG-Cdh1 IPs were compared to other APC/C substrate discovery efforts (47,82). Singh et al. identified “clusters” of proteins whose levels changed at mitotic exit. For each cluster, they reported a top percentile, and for the clusters that most accurately revealed APC/C substrates (1, 2, and 3), we compile their data in Supplemental Table 3 in terms of which KEN-dependent substrates were identified. Their data from Cluster 1, which identified the most KEN-containing APC/C substrates, is shown in Figure 1C. Lafranchi et al. rank ordered proteins based on the degree of change from mitosis to G1, analyzed by proteomics. We curated their data to identify the cut-off point where the last KEN-dependent APC/C substrate was identified among their rank ordered list. Since they provided no cut-off point, the data comparison in Figure 1C represents the best estimate of their ability to capture APC/C substrates.

### Cell Culture

HeLa, HeLa S3, U2OS, RPE-1, and HCT116 cells were grown in 10% FBS with high glucose DMEM without antibiotics. Cell culturing utilized standard laboratory practices whereby cells were grown and incubated at 37°C containing 5% CO_2_. Frozen cell stocks were stored under liquid nitrogen in 10% DMSO/90% FBS.

GFP-UHRF1^WT^ and GFP-UHRF1^KEN:AAA^ stable overexpression cells were generated by transducing HeLa S3, U2OS, and RPE-1-hTERT cell lines with pHAGE-GFP lentivirus that had been produced in HEK293T cells. Infections were performed in the presence of 8µg/mL polybrene for 48 hours prior to antibiotic selection. Cells were selected for 5-7 days with 8ug/mL (HeLa S3 and U2OS) or 10ug/mL (RPE-1) Blasticidin. Lentiviral particles were produced by transfecting HEK293T cells with Tet, VSVg, Gag/pol, and Rev viral packaging vectors together with the pHAGE-GFP lentiviral vectors using *Trans*IT® MIRUS. Viral particles were collected 48 and 72 hours after transfection and stored at −80°C prior to transduction.

To generate the rescue cell lines, the U2OS and HeLa S3 stable GFP-UHRF1^WT^ and GFP-UHRF1^KEN:AAA^ expression cell lines were transduced with previously described and validated pLKO.1 lentiviral vectors encoding either shControl or 3’UTR targeting shUHRF1 (66), using 8ug/mL polybrene to aid infection. After 48 hours, cells were selected with 2µg/mL Puromycin for 3-5 days. Viral particles were produced by transfecting HEK293T cells with the pLKO.1 constructs and psPAX2 and pMD2.G packaging vectors using *Trans*IT® MIRUS (cat no. MIR 2700), collecting after 48 and 72 hours as mentioned previously.

Mitotic block was induced by treating 25% confluent HeLa S3 cells with 2mM thymidine for 24 hours. After washing the plates three-four times with warm media and incubating in drug-free media for 3-4 hours, cells were treated with 100 ng/mL nocodazole for 10-11 hours prior to harvesting by mitotic shake-off. Samples were washed three or four times with warm media, counted, and re-plated for indicated timepoints.

To synchronize cells in G1/S, HeLa S3 were plated at 20% confluency prior to addition of 2mM thymidine. After 16 hours, cells were washed three times with warm media and left to incubate for 8 hours before the second block in 2mM thymidine for another 16 hours. Cells were washed three times in warm media and collected at specific timepoints as they progress through the cell cycle.

To transiently inactivate the APC/C, HCT116 or U2OS cells were treated with 15µM proTAME (Thermo Fisher cat no. I-440-01M), a pan-APC/C inhibitor (83), for 90 minutes prior to harvest and immunoblotting. Cells had been released from nocodazole-induced mitotic block for 90 minutes in drug-free media prior to addition of drug.

### *In vivo* APC/C Activation assay

70-80% confluent U2OS cells were transfected with the indicated plasmids for 24 hours and then exchanged into fresh media. Alternatively, untransfected cells were used to analyze endogenous proteins. After an eight-hour incubation in fresh media following transfection, cells were treated with 250ng/mL nocodazole for 16 hours. Mitotic cells were isolated by shake-off, washed once in pre-warmed media, counted, and divided equally among 15mL conical tubes. Cells in suspension were treated with DMSO, RO-3306 (10 µM), Roscovitine (10 µM), or MG-132 (20µM) for the indicated amount of time at 37°C. Identical volumes of cells were removed from cell suspensions by pipetting, isolated by centrifugation, and frozen at −20°C prior to processing for immunoblot.

### Molecular Biology

Plasmid transfection of HEK293T, U2OS, and HCT116 was performed with either MIRUS or PolyJet (cat no. SL100688) at 1:3 or 1:4 DNA:plasmid ratio on cells with 50-60% confluency. After 24 hours, the media was changed, and cells were expanded to larger dishes as needed. Samples were collected 24-48 hours after siRNA transfection was performed using a 1:3 ratio of RNAi oligonucleotide to RNAiMAX (cat no. 13778-030). UHRF1 was cloned into the indicated lentiviral vectors mentioned previously using standard gateway recombination cloning. Other APC/C substrates tested for binding to Cdh1 or degradation in the APC/C activation assay were obtained from either the ORFeome collection and cloned into the indicated vectors using gateway recombination cloning or from addgene (see supplemental table) (84).

### Cell lysis and immunoblotting

Cells were lysed on ice for 20 minutes in Phosphatase Lysis buffer (50 mM NaH_2_PO_4_, 150 mM NaCl, 1% Tween-20, 5% Glycerol, pH 8.0, filtered) or NETN (20 mM Tris pH 8.0, 100 mM NaCl, 0.5 mM EDTA, 0.5% NP40) supplemented with 10µg/mL each of aprotonin, pepstatin A, and leupeptin, 1mM sodium orthovanadate, 1mM NaF, and 1mM AEBSF. Following incubation on ice, cell lysates were centrifuged at (20,000 x g) in a benchtop microcentrifuge at 4°C for 20 minutes. Protein concentration was estimated by BCA assay (Thermofisher cat no. PI-23227) according to manufacturer’s protocol. Cell extracts were diluted with SDS-PAGE Gel Loading Buffer (Laemmli Buffer) prior to analysis by SDS-PAGE. Typically, 20-40 µg of protein were loaded on SDS gels (either BioRad 4-12% Bis-Tris or homemade SDS-PAGE gels) and separated at 140-200V for approximately 1 hour. Proteins were transferred by wet-transfer methods onto nitrocellulose membrane, typically at 100V for 1 hour or 10-17V overnight at 4°C. Nitrocellulose membranes were then incubated with TBST (137mM NaCl, 2.7mM KCl, 25mM Tris pH 7.4, 0.5% Tween-20) supplemented with either 5% bovine serum albumin or non-fat dry milk for at least one hour or overnight at 4°C. Blocked membranes were incubated overnight with primary antibodies at 4°C, washed in TBST, incubated in appropriate secondary antibodies for 1 hour at room temperature, and then developed by chemiluminescence using Pierce ECL (ThermoFisher) or Clarity ECL (Bio-Rad). See reagent list in supplement for detailed primary and secondary antibody information.

### Immunoprecipitation

For co-immunoprecipitation (coIP) experiments, cells were lysed in NETN for 20 minutes on ice and then centrifuged in a benchtop centrifuge on maximum speed (20,000 x g) for 20 minutes at 4°C, prior to determining protein concentration by either Bradford or BCA assay.

A master mix of 1-2 mg/mL protein concentration was calculated, 10% of which was retained as input while the remaining 90% was used for coIP. Prior to coIP, antibody coated beads were prewashed with 1X TBST three times prior to incubation with lysis buffer. Cell lysates were also pre-cleared by incubation with the same volume of empty Protein A/G agarose beads. Clarified cell lysates were immunoprecipitated for 2-4 hr at 4°C with 25-50uL of EzView M2- or Myc-antibody beads (F2426-1ML or E6654-1ML). After coIP, beads were pelleted at low speed centrifugation, washed twice with wash buffer, and one time with lysis buffer to remove unbound proteins. Buffers were removed from beads using a 27 gauge needle to avoid the aspiration of beads between washes. Washed beads were resuspended in 2X SDS-PAGE Gel Loading Buffer (Laemmli Buffer) and boiled 5-10 minutes at 95°C. Samples were removed from the beads using a 27-gauge needle to avoid the aspiration of beads after boiling. Typically, 20µL of coIP was loaded alongside 1% of the input volume. Samples were analyzed by immunoblotting as described.

### Protein Purification

Substrates for *in vitro* ubiquitylation assays were expressed as N-terminal GST-TEV-fusion (TTF2) or His-MBP-TEV-fusions (FL-UHRF1^WT^, LPS-UHRF1^WT^, FL-UHRF1^KEN:AAA^, LPS-UHRF1^KEN:AAA^) in BL21 (DE3) codon plus RIL cells. TTF2 was purified by glutathione-affinity chromatography, treated with TEV protease to liberate GST, and further purified by ion exchange chromatography. UHRF1 wild-type and variants were purified by amylose-affinity chromatography, treated with TEV, and followed by ion exchange chromatography. Fluorescently labeled substrates were generated by incubating 1 µM Sortase, 20x 5-carboxyfluorescein (5-FAM)-PEG-LPETGG peptide, and substrates in 10 mM HEPES pH 8, 50 mM NaCl, and 10 mM CaCl_2_. After 2 hours of incubation at 4*°*C, reactions were stopped by removing the His_6_-tagged Sortase by nickel affinity chromatography. Then, excess 5-FAM-LPETGG was removed by size exclusion chromatography.

Expression and purification of UBA1, UBE2C, UBE2S, recombinant APC/C and pE-APC/C, Cdh1, Cdc20, Emi1, ubiquitin, and methylated ubiquitin were performed as described previously in Brown et al. 2016 (85–89).

### APC/C Ubiquitylation assays

Qualitative assays to monitor APC/C-dependent ubiquitylation were performed as previously described (89). In brief, reactions were mixed on ice, equilibrated to room temperature before the reactions are initiated with Ub or meUb, and quenched at the indicated time points with SDS. TTF2 ubiquitylation was monitored by mixing 100 nM APC/C, 1 µM Cdh1, 5 µM UBE2C, 5 µM UBE2S (when indicated), 1 µM UBA1, 5 µM TTF2, 5 mM Mg-ATP, and 150 µM Ub or meUb (Fig. S2). Ubiquitylation of UHRF1 wild-type or its variants by APC/C were performed with 100 nM APC/C or pE-APC/C, 1 µM Cdh1 or Cdc20, 0.4 µM UBE2C, 0.4 µM UBE2S (when indicated), 1 µM UBA1, 0.4 µM UHRF1, 5 mM Mg-ATP, and Ub or meUb (Fig. 4 and Fig. S4). Following SDS-PAGE, ubiquitylation products of the fluorescently labeled substrates were resolved by SDS-PAGE and imaged with the Amersham Typhoon 5.

### Flow cytometry cell cycle analysis

HeLa S3 GFP-UHRF1^WT^ and GFP-UHRF1^KEN:AAA^ (shUHRF1) cells were synchronized in mitosis by sequential thymidine-nocodazole treatment as described above, using 2mM thymidine and 100ng/mL nocodazole. After release, cells were pulsed with 10µM EdU thirty minutes prior to collection at specific timepoints. After counting the cells, 2 million cells were retained for Western blotting (WB) analysis and 1 million cells were fixed for flow cytometry. For WB, cells we pelleted and washed once with cold PBS prior to freezing at −20°C. For flow cytometry, cells were fixed in 4% formaldehyde/PBS for 15 minutes at room temperature. Cells were pelleted and resuspended in 1% BSA/PBS and stored overnight at 4°C. The next day, cells were pelleted and resuspended in 1% BSA/PBS/0.5% Triton X-100 for 15 minutes at room temperature. Cells were pelleted, resuspended with labelling solution (100mM ascorbic acid, 1mM CuSO_4_, 2µM Alexa Fluor 488 azide in PBS), and incubated for thirty minutes in the dark at room temperature. After addition of 1% BSA/PBS/0.5% Triton X-100, cells were pelleted and stained with 1µg/mL DAPI in 1% BSA/PBS/0.5% Triton X-100 for one hour in the dark at room temperature. Flow cytometry was performed on an Attune™ Nxt Flow Cytometer (Thermo Fisher Scientific). Channel BL1 was used for Azide 488 dye. Channel VL1 was used for DAPI dye. Following acquisition, data were analyzed using FlowJo software.

### Immunofluorescence imaging

HeLa cells were plated on poly-L-lysine-coated #1.5 coverslips. Next day, cells were treated with siRNA (control siFF and siUHRF1) and RNAi Max according to manufacturer’s protocol (Invitrogen). After 48 hours of siRNA treatments, cells were fixed in 3% paraformaldehyde in PHEM buffer (60 mM PIPES, 25 mM HEPES, 10 mM EGTA, 2 mM MgCl_2_, pH 7.0) for 15 minutes at 37 °C. Then, cells were washed with PHEM buffer and permeabilized using 0.5% of Nonidet P-40 in PHEM buffer for 15 minutes at room temperature. Cells were washed and then blocked with 5% BSA in PHEM. Primary antibodies used were: α-CENP-C (MBL:1:1000) as a kinetochore marker and α-tubulin (Sigma: 1:500). Samples were incubated in primary antibody solution for 1 hour at 37 °C. All fluorescently labeled secondary antibodies (anti-mouse Alexa 488, anti-guinea pig 564) were diluted 1:200 dilution, and cells were incubated for 1 hour at 37 °C. DNA was counterstained with DAPI for 15 minutes at room temperature after washing out secondary antibodies. All samples were mounted onto glass slides in Prolong Gold antifade (Invitrogen). For image acquisition, three-dimensional stacked images were obtained sequentially at 200 nm steps along the z axis through the cell using MetaMorph 7.8 software (Molecular Devices) and a Nikon Ti-inverted microscope equipped with the spinning disc confocal head (Yokogawa), the Orca-ER cooled CCD camera (Nikon), and an ×100/1.4 NA PlanApo objective (Nikon).

### Genomic DNA isolation for methylation analysis

Genomic DNA was isolated from Parental U2OS cells and U2OS cells overexpressing either GFP-UHRF1^WT^ or GFP-UHRF1^KEN:AAA^. All samples groups were processed in biological triplicates. Briefly, cells were lysed overnight at 37°C in 2 mL of TE-SDS buffer (10 mM Tris-HCl pH 8.0, 0.1 mM EDTA, 0.5% SDS), supplemented with 100 μl of 20 mg/ml proteinase K. DNA was purified by phenol:chloroform extraction in three phases: (1) 100% phenol, (2) phenol:chloroform:isoamyl alcohol (25:24:1), and (3) chloroform:isoamyl alcohol (24:1). For each phase, the aqueous layer was combined with the organic layer in a 1:1 ratio. Samples were quickly shaken, allowed to sit on ice for approximately 5 minutes, and then separated by centrifugation at 1,693 RCF for 5 minutes at 4°C. The top aqueous layer was then transferred to a new tube for the next organic phase. Following extraction, DNA was precipitated with 1/10 volume 3M sodium acetate pH 4.8 and 2.5 volumes 100% ethanol and stored overnight at −20°C. Precipitated DNA was pelleted by centrifugation at 17,090 RCF for 30 minutes at 4°C. The pelleted DNA was washed twice with 70% ethanol, allowed to dry for 15 minutes, and resuspended in TE buffer (10 mM Tris-HCl pH 8.0, 0.1 mM EDTA). Samples were then treated with 1 mg/ml RNAse A at 37°C for 30 minutes and then re-purified by ethanol precipitation as described above.

### Infinium Methylation EPIC BeadChip (EPIC array)

Genomic DNA was quantified by High Sensitivity Qubit Fluorometric Quantification (Invitrogen), and 1.5 ug of genomic DNA was submitted to the Van Andel Institute Genomics Core for quality control analysis, bisulfite conversion, and DNA methylation quantification using the Infinium Methylation EPIC BeadChIP (Illumina) processed on an Illumina iScan system following the manufacturer’s standard protocol (67,68).

### EPIC array data processing

All analyses were conducted in the R statistical software (Version 3.6.1) (R Core Team). R script for data processing and analysis is available in Supplemental Code File 1.

Raw IDAT files for each sample were processed using the Bioconductor package “SeSAMe” (Version 1.2.0) for extraction of probe signal intensity values, normalization of probe signal intensity values, and calculation of β-values from the normalized probe signal intensity values (90–92). The β-value is the measure of DNA methylation for each individual CpG probe, where a minimum value of 0 indicates a fully unmethylated CpG and a maximum value of 1 indicates a fully methylated CpG in the population. CpG probes with a detection p-value > 0.05 in any one sample were excluded from the analysis.

### Genomic and Replication Timing annotation

CpG probes were mapped to their genomic coordinate (hg38) and were then annotated to their genomic annotation relationship (promoter-TSS, exon, etc.) using HOMER (Version 4.10.3) (93).

Repli-seq data for U2OS cells used for determining CpG probe localization relative to replication timing was generated by Dr. David Gilbert’s lab (Florida State University) as part of the 4D Nucleome project (Experiment #4DNEXWNB33S2)(69). Genomic regions were considered early-replicating if the replication timing value was > 0 and late-replicating if < 0. CpG probes were annotated for replication timing domains by intersecting the Repli-seq genomic coordinates with CpG probe coordinates using BEDTools (Version 2.16.2) (94).

### Identification of differentially methylated CpG probes

The Bioconductor package “limma” (Version 3.40.6) was used to determine differential methylation among sample groups and perform multidimensional scaling (MDS) analysis (95–97). For statistical testing of significance, β-values were logit transformed to M-values: 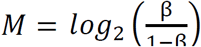. M-values were then used for standard limma workflow contrasts to determine differential methylation of U2OS GFP-UHRF1^WT^ or GFP-UHRF1^KEN:AAA^ overexpression to Parental U2OS cells (98,99). CpG probes with an adjusted p-value ≤ 0.05 were considered significant, and log fold-change of M-value was used to determine hypermethylation (logFC > 0) or hypomethylation (logFC < 0) relative to U2OS parental cells.

### Enrichment Bias Calculation and Hypergeometic Distribution Testing

Enrichment Bias Calculations were done by first determining the following values for each feature (e.g. Genomic Annotation, Replication Timing):

*q* = Number of CpGs that are differentially methylated in feature (e.g. exon)

*m* = Total number of CpGs on the EPIC array that match feature (e.g. exon)

*n* = Total number CpGs on the EPIC array that do not match feature (e.g. everything that is not an exon)

*k* = Total number of all differentially methylated CpGs

Next, the expected number of CpGs that would be differentially methylated in that feature by random chance was determined with the following equation:

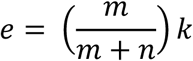

Finally, percent enrichment bias was calculated with the following equation:

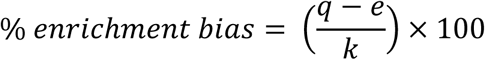

Where positive or negative enrichment values indicate more or less enrichment for a feature than would be expected by random chance, respectively.

Hypergeometric distribution testing for determining significance of enrichment bias was performed using the phyper() function in R with the following values: *q,m,n,k*.

### Data access

EPIC array data can be found under GEO Accession # GSE137913.

To review GEO accession GSE137913:

Go to https://www.ncbi.nlm.nih.gov/geo/query/acc.cgi?acc=GSE137913

The following secure token has been created to allow review of record GSE137913 while it remains in private status: eletaomyfnqrlun

### Signature evaluation in TCGA BRCA samples

Upper quartile normalized RSEM gene expression data for TCGA BRCA (n=1201) was downloaded from the GDC legacy archive (https://portal.gdc.cancer.gov). The data was log2 transformed and median centered. To determine the per sample UB signature score, the samples were ranked by the median expression of the 145 UB gene signature. Sample were then divided at the median and grouped as high or low based on rank. Copy number burden, aneuploidy, and homologous recombination deficiency data were extracted from Thorsoon et. al. (100) and plotted by UB signature group and PAM50 subtype (101). Significance was calculated by t-test. The CIN70 score was determined as previously described in Fan et. al. (102). The CIN70 was plotted against the UB, colored by PAM50 subtype, and r^2^ and Pearson correlation were calculated. All analysis were performed in R (v3.5.2).

### Cdh1 pulldown for analysis of interactors by mass spectrometry

FLAG-tagged Cdh1 was expressed in HEK293T cells for 24 hours by transient transfection. Transfections were performed on 150 mm dishes (8 per condition) using Mirus TransIT®-LT1 Transfection Reagent (Mirus Bio) and Lipofectamine 2000 (Life Technologies). Cells were treated with MG-132 (10 μM for 4 hours) in culture prior to lysis, dislodged by trypsinization, washed with PBS, and lysed in NETN supplemented with 2 μg/ml pepstatin, 2 μg/ml apoprotinin, 10 μg/ml leupeptin, 1 mM AEBSF (4-[2 Aminoethyl] benzenesulfonyl fluoride), 1 mM Na_3_VO_4_, and 1 mM NaF on ice for 20 minutes. Cell lysates were then clarified by centrifugation at 15,000 rpm for 15 minutes.

Anti-FLAG M2 agarose (Sigma, catalog no. F2426) was used for precipitation (6 hours at 4°C). The beads were washed with NETN three times and eluted twice with 150 µl of 0.1 M Glycine-HCl, pH 2.3 and then neutralized with Tris 1M (pH 10.0). The total eluted protein was reduced (5 mM DTT) and alkylated using iodoacetamide (1.25 mM) for 30 minutes in the dark. The resultant protein was then digested overnight with sequencing grade trypsin (Promega). The trypsin: protein ratio was maintained at 1:100. Total peptides were purified on Pierce C18 spin columns (Cat 89870) using the manufacturer’s protocol. Peptides were eluted using 70% acetonitrile and 0.1% TFA solution in 50 μl volumes twice, dried on a SpeedVac at room temperature, and processed by mass spectrometry proteomic analysis.

### Mass Spectrometry

Peptides were separated by reversed-phase nano-high-performance liquid chromatography using a nanoAquity UPLC system (Waters Corp.). Peptides were first trapped in a 2 cm trapping column (Acclaim® PepMap 100, C18 beads of 3.0 μm particle size, 100 Å pore size) and a 25 cm EASY-spray analytical column (75 μm inner diameter, C18 beads of 2.0 μm particle size, 100 Å pore size) at 35°C. The flow rate was 250 nL/minute over a gradient of 1% buffer B (0.1% formic acid in acetonitrile) to 30% buffer B in 150 minutes, and an in-line Orbitrap Elite mass spectrometer (Thermo Scientific) performed mass spectral analysis. The ion source was operated at 2.6 kV with the ion transfer tube temperature set at 300°C. A full MS scan (300–2000 m/z) was acquired in Orbitrap with a 120,000 resolution setting, and data-dependent MS2 spectra were acquired in the linear ion trap by collision-induced dissociation using a 2.0 m/z wide isolation window on the 15 most intense ions. Precursor ions were selected based on charge states (+2, +3) and intensity thresholds (above 1e5) from the full scan; dynamic exclusion (one repeat during 30 seconds, a 60 seconds exclusion time window) was also used. The polysiloxane lock mass of 445.120030 was used throughout spectral acquisition.

Raw mass spectrometry data files were searched using SorcererTM-SEQUEST® (build 5.0.1, Sage N Research), the Transproteomic Pipeline (TPP v4.7.1), and Scaffold (v4.4.1.1) with the UniProtKB/Swiss-Prot human canonical sequence database (20,263 entries; release 07/2013). The search parameters used were a precursor mass between 400 and 4500 amu, zero missed cleavages, a precursor ion tolerance of 3 amu, accurate mass binning within PeptideProphet, fully tryptic digestion, a static carbamidomethyl cysteine modification (+57.021465), variable methionine oxidation (+15.99492), and variable serine, threonine and tyrosine (STY) phosphorylation (79.966331). A 1% protein-level FDR was determined by Scaffold.

## Acknowledgements

We thank Brian Strahl (UNC) and members of the Emanuele and Brown labs for helpful discussions throughout this project. We thank Marie Adams and Julie Koeman from the Van Andel Institute Genomics Core for their assistance with Illumina Infinium MethylationEPIC BeadChip processing. We acknowledge the UNC Flow Cytometry Core Facility (supported in part by P30 CA016086 Cancer Center Core Support Grant to the UNC Lineberger Comprehensive Cancer Center). The Emanuele lab (JLK, MJE, XW, TB, and RC) is supported by the UNC University Cancer Research Fund (UCRF), the National Institutes of Health (R01GM120309), the America Cancer Society (Research Scholar Grant; RSG-18-220-01-TBG). The Hoadley lab (KAH and JD) is supported by a Komen Career Catalyst Award (CCR16376756). The Brown lab (RCMC, DLB, and NGB) is supported by UCRF and National Institutes of Health (R35GM128855). The Suzuki lab is supported by startup funds from the University of Wisconsin-Madison School of Medicine, Graduate School, McArdle Laboratory for Cancer Research, the UW Carbone Cancer Center, and by the New Investigator Program grant from Wisconsin Partnership Program (WPP). The Rothbart lab (SBR and RLT) are supported by grants from the NIH (R35GM124736) and the American Cancer Society – Michigan Cancer Research Fund (PF-16-245-01-DMC).

## Author Contributions

JLK performed most cell biological experiments. RCMC and DLB purified recombinant proteins and performed *in vitro* ubiquitylation assays. TB and XW performed cell-based assays. RC, FY, and MBM performed proteomics experiments. JSD and KAH performed bioinformatic analysis of TCGA datasets. AS performed imaging analysis of mitotic cells. SBR and RLT performed DNA methylation analysis. JSH provided UHRF1 constructs and technical advice on protein purification. MJE conceived of the computational screen and analyzed data. NGB and MJE designed experiments. JLK, RCMC, RLT, SBR, NGB, and MJE wrote and revised the manuscript.

## Supplemental Figure Legends

**Fig. S1.**
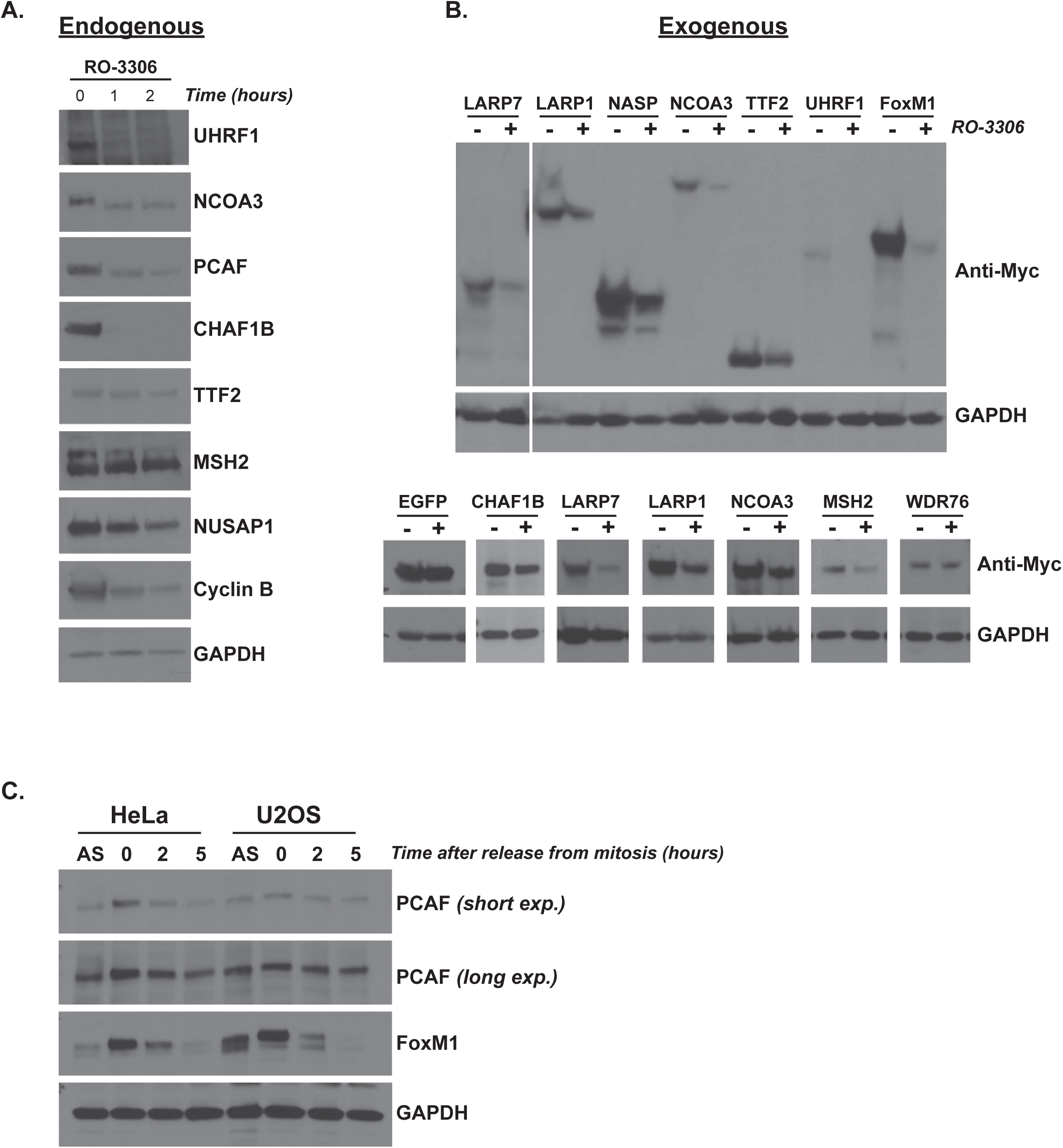
Analysis of putative APC/C substrates. (A) U2OS cells were arrested in mitosis with nocodazole, collected by shake-off, treated with the CDK1 inhibitor RO-3306, and harvested for immunoblot at the indicated timepoints. Cyclin B and NUSAP1 serve as positive APC/C controls. (B) U2OS cells were transiently transfected with the indicated plasmids, arrested in mitosis with nocodazole, collected by shake-off, treated with the CDK1 inhibitor RO-3306, and harvested for immunoblot after 2 hr. FoxM1 serves as a positive control for APC/C activation. (C) HeLa and U2OS cells were synchronized in mitosis by nocodazole and released by mitotic shake-off. Timepoints were collected as shown and analyzed by immunoblot. FoxM1 serves as positive APC/C control that is degraded at M/G1 phases.

**Fig. S2.**
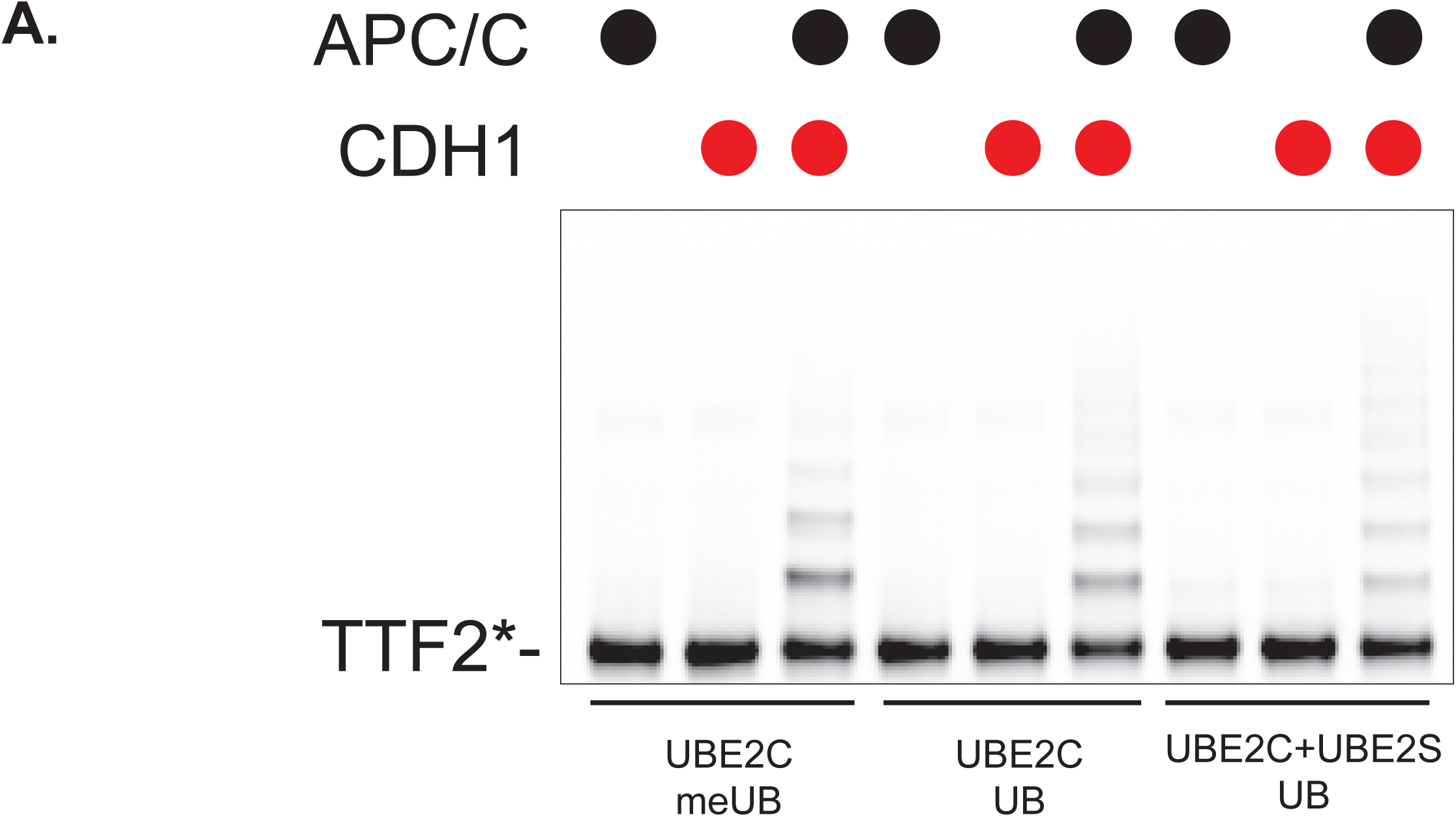
TTF2 is ubiquitylated by APC/C *in vitro*. (A) Ubiquitylation reactions of TTF2* by UBE2C using methylated Ub or wild-type Ub (lanes 1-6) in combination with APC/C^Cdh1^, APC/C alone, or Cdh1 alone. Ubiquitylation reactions of TTF2* by both E2s, UBE2C and UBE2S, (lanes 7-9) in combination with APC/C^Cdh1^, APC/C alone, or Cdh1 alone. Ubiquitylation was detected by fluorescence scanning at 60 minute timepoints.

**Fig. S3.**
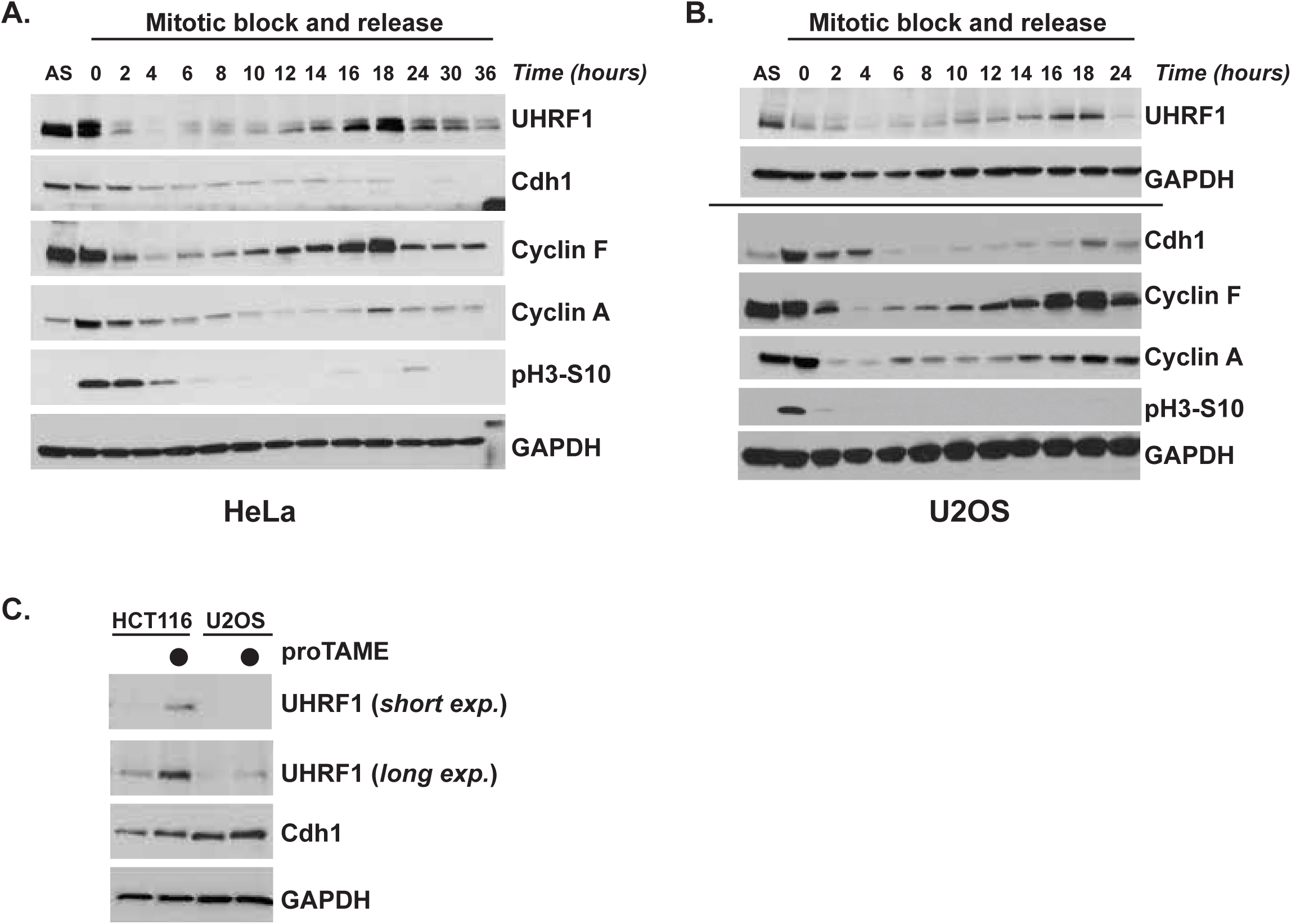
UHRF1 protein levels are cell cycle regulated and sensitive to APC/C inhibition with the small-molecule inhibitor proTAME. (A) HeLa cells were synchronized in mitosis, collected by shake-off, released into the cell cycle, and analyzed by immunoblot at the indicated timepoints. (B) U2OS cells were synchronized in mitosis, collected by shake-off, released into the cell cycle, and analyzed by immunoblot at the indicated timepoints. Line indicates samples that were run on separate gels, with appropriate corresponding loading controls for each gel. (C) HCT116 and U2OS cells were released into G1 from a mitotic block for 1.5hr and then were subsequently treated with proTAME for 1.5 hr. Endogenous UHRF1 and Cdh1 were analyzed by immunoblot.

**Fig. S4.**
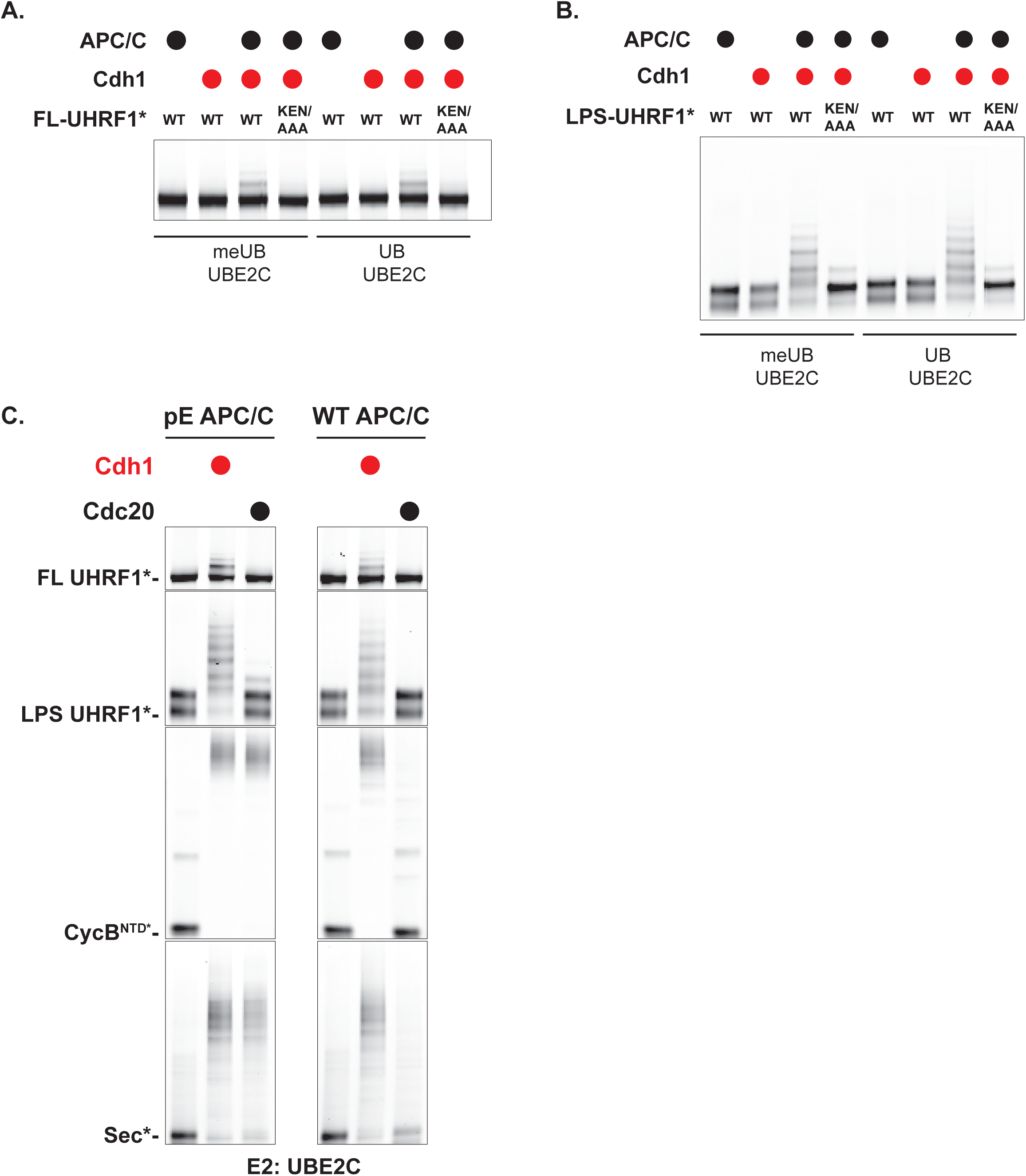
UHRF1 ubiquitylation by APC/C. (A) Ubiquitylation reactions of FL-UHRF1* by UBE2C with either methylated Ub or wild-type Ub. Reactions were performed using UHRF1^WT^ or a variant harboring alanine substitution in the KEN-box (KEN:AAA). KEN degron motif mutants in UHRF1 are shown in lanes 4 and 8. Ubiquitylation was detected by fluorescence scanning at 30 minute timepoints. (B) Ubiquitylation reactions of LPS-UHRF1* by UBE2C with either methylated Ub or wild-type Ub. Reactions were performed using UHRF1^WT^ or a variant harboring alanine substitution in the KEN-box (KEN:AAA). KEN degron motif mutants in UHRF1 are shown in lanes 4 and 8. Ubiquitylation was detected by fluorescence scanning at 30 minute timepoints. (C) Ubiquitylation reactions of FL-UHRF1* and LPS-UHRF1* are exclusive to Cdh1 as the coactivator. Ubiquitylation reactions were performed using wild-type APC/C^Cdh1^ which can only utilize Cdh1, but not Cdc20, as well as pE-APC/C^Cdh1^, which mimics the APC/C phosphorylated state and can therefore use either Cdc20 or Cdh1. In parallel, we analyzed ubiquitylation of CycB^NTD*^ and Securin*, which can be ubiquitylated by both APC/C^Cdc20^ and APC/C^Cdh1^.

**Fig. S5.**
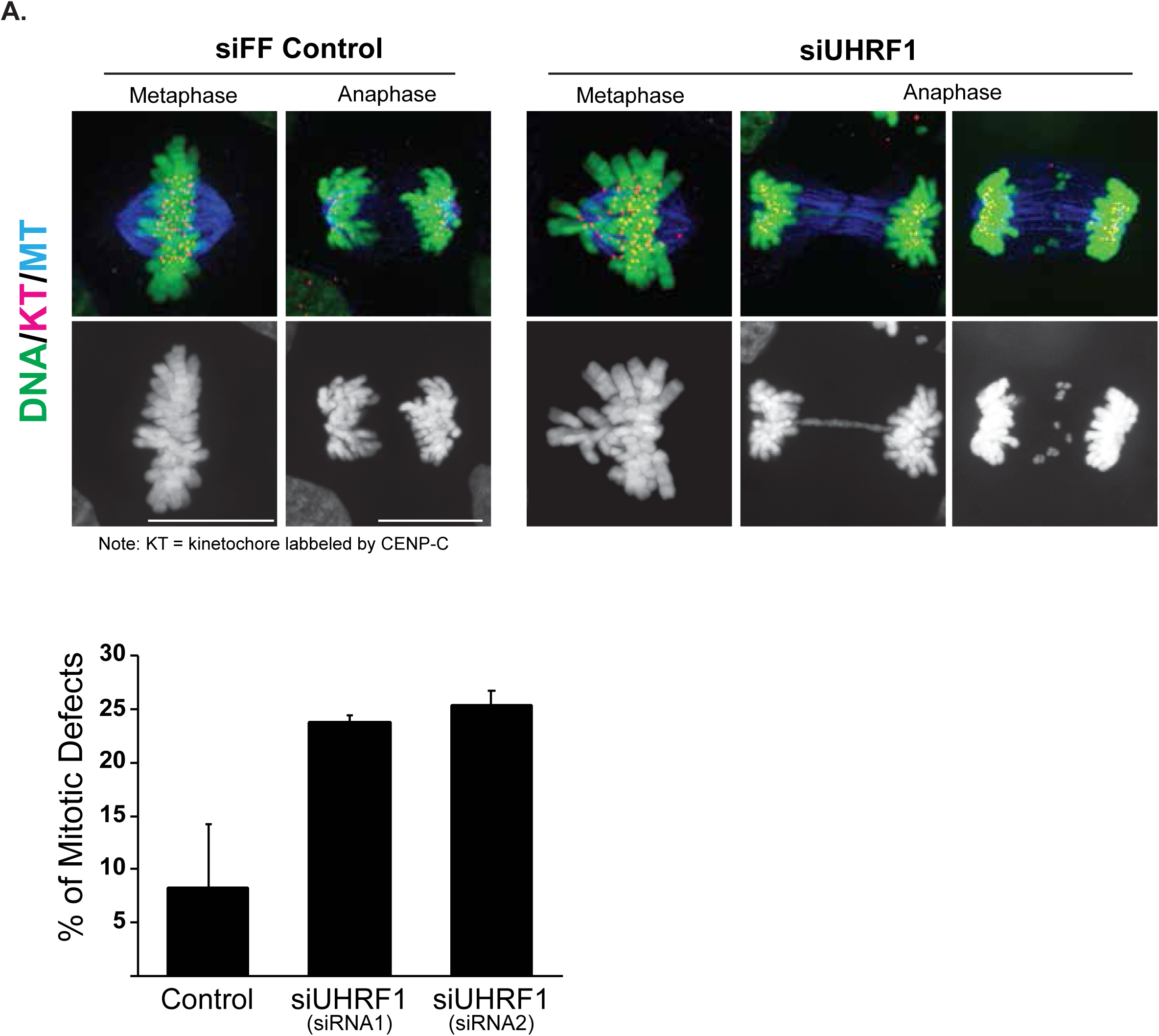
UHRF1 depletion impairs chromosome alignment. (A) HCT116 cells were depleted of UHRF1 using two independent siRNA oligonucleotides. Cells were fixed and stained with antibodies to the kinetochore protein CENP-C and microtubules.

**Fig. S6.**
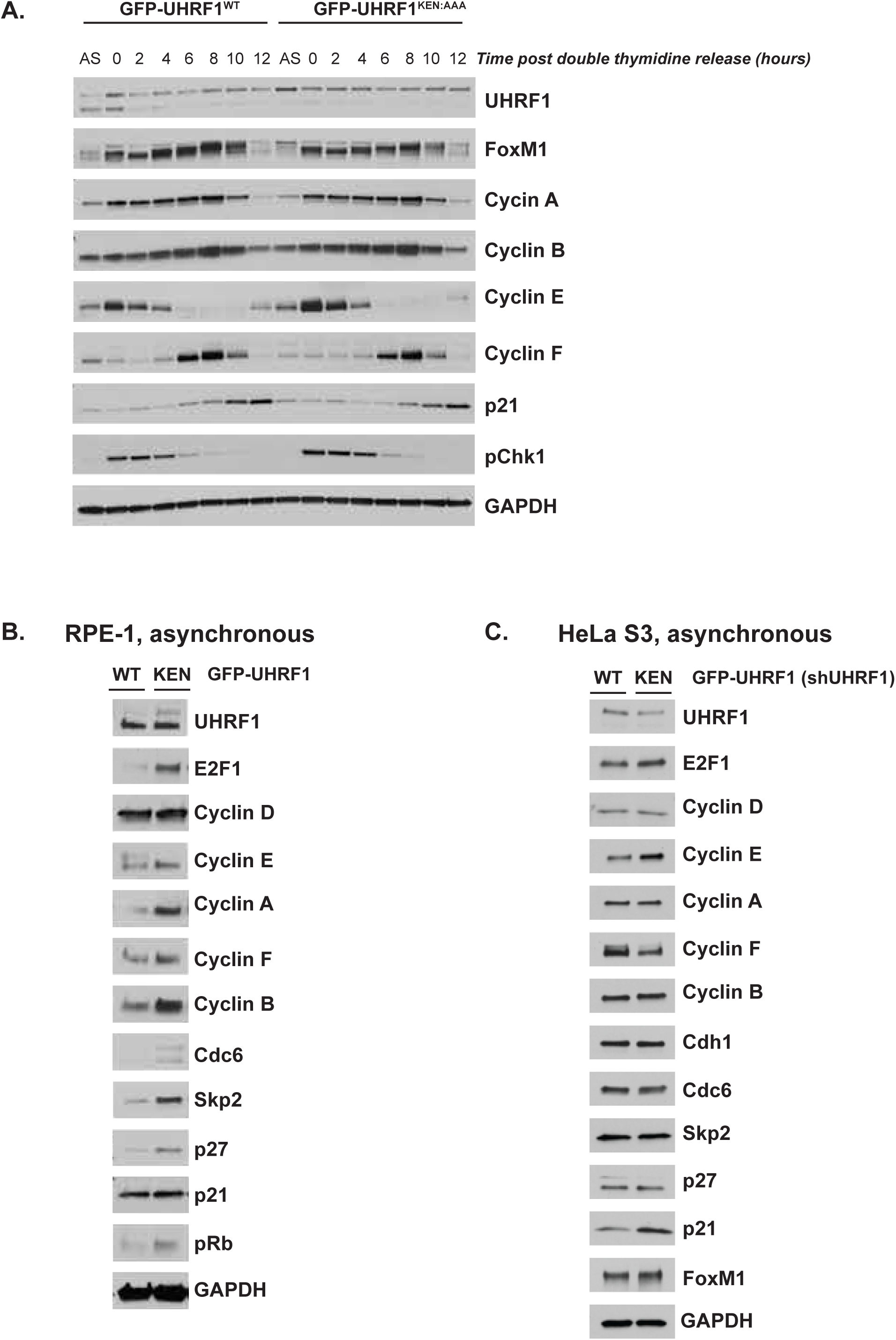
Progression through S/G2 phases in cells expressing non-degradable UHRF1. (A) HeLa S3 cells stably expressing GFP-UHRF1^WT^ or GFP-UHRF1^KEN:AAA^ were synchronized at G1/S by double thymidine block, released in the cell cycle, and analyzed by immunoblot at the indicated time points. Cells progressed through S/G2 phases with minimal differences except for an increase in cyclin E levels. (B) Asynchronous RPE-1 cells stably expressing GFP-UHRF1^WT^ or GFP-UHRF1^KEN:AAA^ were harvested for immunoblotting for cell cycle markers as shown. (C) Asynchronous HeLa S3 cells stably expressing GFP-UHRF1^WT^ or GFP-UHRF1^KEN:AAA^ along with 3’UTR targeting shUHRF1 were harvested for immunoblotting for cell cycle markers as shown.

**Fig. S7.**
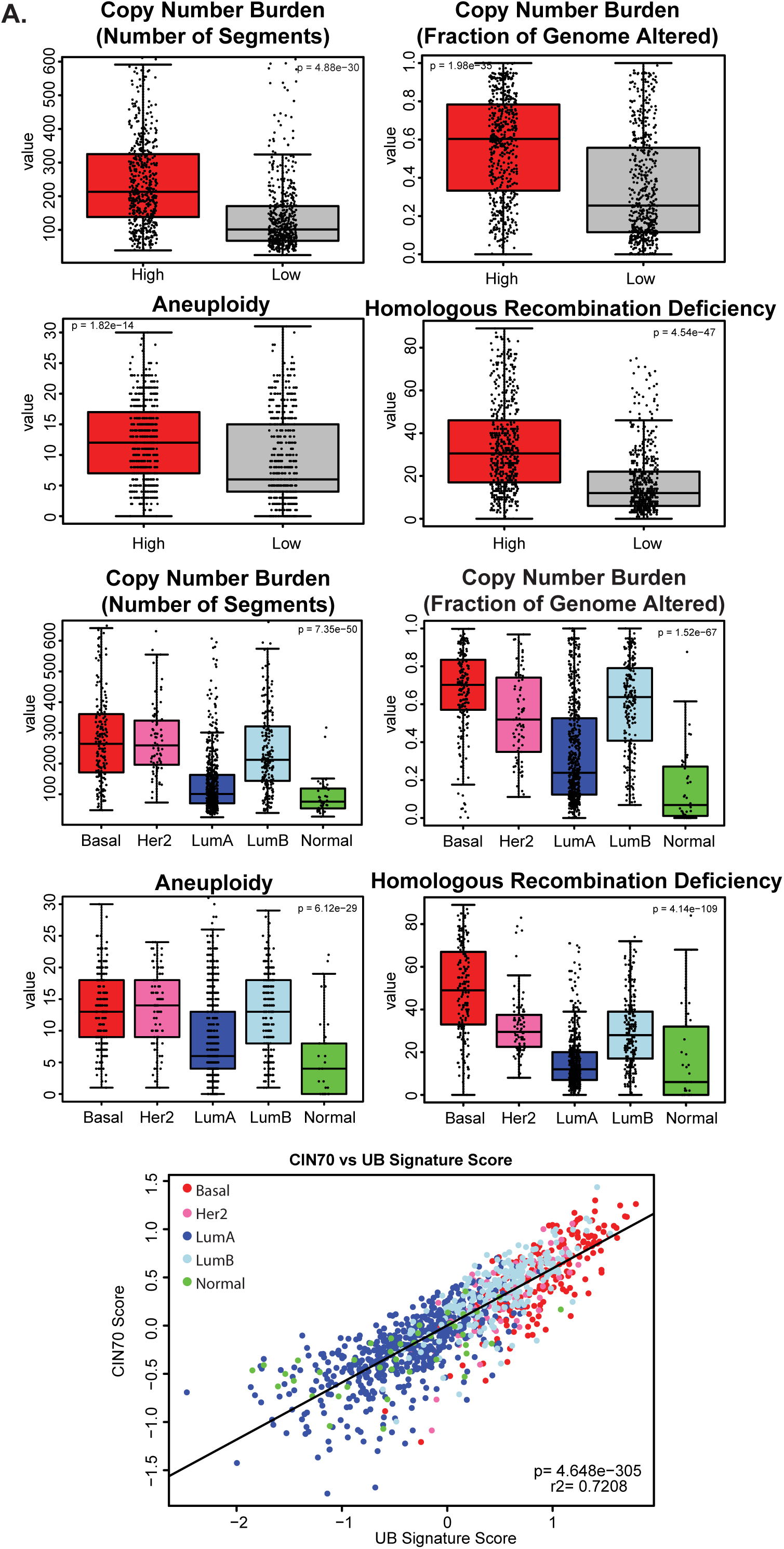
A 145 gene signature derived from KEN-containing proteins which have cell cycle dependent gene transcription is associated with makers of chromosome instability in breast cancer. (A) TCGA BRCA samples (n=1201) were assigned to High or Low based on the ranked median value of the 145 gene signature score. Samples were then plotted for the given genomic feature based on Thorsson et. al. by both gene signature group and PAM50 subtype. Significant was determined by t-test or ANOVA where appropriate. The median 145 gene signature score was plotted against the chromosome instability score (CIN70) (r2=0.72, Pearson correlation p<0.001). Colors indicate PAM50 subtypes.

